# Integrated proteomics reveals the landscape of autophagy degradation in human neurons and autophagy receptors regulating neuronal activity

**DOI:** 10.1101/2022.12.04.519029

**Authors:** Xiaoting Zhou, You-Kyung Lee, Xianting Li, Henry Kim, Carlos Sanchez-Priego, Xian Han, Haiyan Tan, Suiping Zhou, Yingxue Fu, Kerry Purtell, Qian Wang, Gay Holstein, Beisha Tang, Junmin Peng, Nan Yang, Zhenyu Yue

**Affiliations:** Department of Neurology, The Friedman Brain Institute, Icahn School of Medicine at Mount Sinai, New York, NY 10029.; Department of Neuroscience, The Friedman Brain Institute, Icahn School of Medicine at Mount Sinai, New York, NY 10029.; Department of Geriatrics, Xiangya Hospital, Central South University, Changsha, Hunan, 410008, China.; Nash Family Department of Neuroscience, Friedman Brain Institute, Icahn School of Medicine at Mount Sinai, New York, NY, 10029, USA.; Black Family Stem Cell Institute, Icahn School of Medicine at Mount Sinai, New York, NY 10029, USA; Department of Structural Biology, St. Jude Children’s Research Hospital, Memphis, TN 38105, USA; Department of Developmental Neurobiology, St. Jude Children’s Research Hospital, Memphis, TN 38105, USA; Integrated Biomedical Sciences Program, University of Tennessee Health Science Center, Memphis, TN 38163, USA; Center for Proteomics and Metabolomics, St. Jude Children’s Research Hospital, Memphis, TN 38105, USA; Department of Neurology, Xiangya Hospital, Central South University, Changsha, Hunan, 410008, China; National Clinical Research Center for Geriatric Disorders, Xiangya Hospital, Central South University, Changsha, Hunan 410008, China

## Abstract

Autophagy is a catabolic and self-degradative process crucial for maintaining cellular homeostasis. Malfunctional autophagy is implicated in neurodevelopmental and neurodegenerative diseases. However, the exact role and targets of autophagy in human neurons remain elusive. Here we reported a systematic investigation of neuronal autophagy targets through integrated proteomics. Deep proteomic profiling of multiple autophagy-deficient lines of human induced neurons, mouse brains, and LC3-interactome uncovers a role of neuronal autophagy in targeting primarily endoplasmic reticulum (ER), mitochondria, endosome, Golgi apparatus, synaptic vesicle (SV) proteins, and cAMP-PKA pathway, for degradation. Tubular ER and specific SV proteins are significant autophagy cargos in the axons. Functional validation identified calumenin as an ER resident autophagy receptor and illuminated a role of autophagy in regulating PKA and neuronal activity through AKAP11-mediated degradation. Our study thus reveals the landscape of autophagy degradation in human neurons and offers molecular insight into mechanisms of neurological disorders linked to autophagy deficiency.

**Highlight:**

1. Integrated proteomics reveals the landscape of autophagy degradation in human neurons
2. Autophagy clears tubular ER and selective ER and synaptic vesicle proteins in neurons
3. Calumenin is an ER resident autophagy receptor
4. Autophagy controls PKA pathway and neuronal activity through autophagy receptor AKAP11

## Introduction

Macroautophagy (herein referred to as autophagy) is a lysosome-dependent degradation pathway. Mammalian cells require autophagy to maintain cellular homeostasis and respond to various stresses and injuries through digestion and recycling. During autophagy, cytoplasmic components are engulfed in a double-membraned vesicle known as the autophagosome, which is shuttled to the lysosome for degradation (Mizushima and Komatsu, 2011). While autophagy degrades intracellular components in the non-selective manner in response to nutrient starvation, autophagy can also selectively degrade cytoplasmic proteins and organelles under basal conditions. More specifically, Atg8-family proteins such as LC3A/B, that are conjugated on the growing autophagosome membrane selectively recognize autophagy cargos (Kabeya et al., 2000). The identification of specific interactions between Atg8-family proteins and autophagy cargos has enabled the utilization of affinity purification and proximity labeling to profile selective autophagy substrates in cancer cell lines(Behrends et al., 2010; Chino et al., 2019; Zellner et al., 2021a).

In contrast to most mammalian cell types, neurons are post-mitotic, long-living cells of the central nervous system (CNS). To maintain longevity and physiological function, neuronal autophagy removes and recycles misfolded proteins, protein aggregates, and injured organelles that would otherwise be detrimental to the cell. In fact, when essential autophagy-related genes (ATGs) *Atg5* and *Atg7* are conditionally deleted in mouse brains, cerebral and cerebellar neurons degenerate, causing motor dysfunctions in mice (Hara et al., 2006; Komatsu et al., 2006). Using mice with disrupted *Atg7* expression in specific neuron types, we previously demonstrated a compartmental function of autophagy in neurons by maintaining axonal homeostasis and preventing axonopathy (Friedman et al., 2012; Komatsu et al., 2007b). Indeed, autophagy biogenesis can occur in the axon and participates in axonal transport to remove dysfunctional organelles and protein aggregates (Goldsmith et al., 2022; Maday and Holzbaur, 2014; Roney et al., 2021). Furthermore, autophagy was shown to regulate synaptic activity (Hernandez et al., 2012; Tang et al., 2014; Vijayan and Verstreken, 2017). The significance of autophagy in the CNS is further underscored in humans, where a recent study has identified biallelic, recessive variants in human *ATG7* that results in impaired autophagic flux and complex neurodevelopmental disorders(Collier et al., 2021). Furthermore, dysfunctional autophagy may contribute to major neurodegenerative diseases (Menzies et al., 2017; Nixon, 2013; Yamamoto and Yue, 2014). However, the physiology of autophagy in human neurons and detailed mechanisms for how its disruption leads to various neurodegenerative diseases remains poorly understood.

To understand the autophagy targets, multiple studies performed systemic analysis to dissect autophagy cargo in non-neuronal cells (An et al., 2019; Chino et al., 2019; Dengjel et al., 2012; Le Guerroué et al., 2017; Mancias et al., 2014; Øverbye et al., 2007; Schmitt et al., 2021; Zellner et al., 2021b). However, the investigation of neuronal autophagy cargo, particularly in humans, remains limited. Profiling the differentially accumulated proteins from *Atg5*-deficient primary mouse neurons has revealed the function of autophagy in regulating axonal endoplasmic reticulum (ER) and luminal Ca2+ stores (Kuijpers et al., 2021; Ordureau et al., 2021). Examination of isolated autophagic vesicles (AVs) from mouse brains using differential centrifugation showed the presence of mitochondrial fragments and synaptic proteins in neuronal autophagosomes (Goldsmith et al., 2022). While these studies have detected cargo candidates of neuronal autophagy, the discrepancy in the findings has yet to clearly define the autophagy targets and mechanism of selectivity.

Herein we present a systemic, comprehensive investigation of human neuronal autophagy cargo by using human pluripotent stem cell (PSC)-induced neurons (iNeurons) through integrated proteomics and functional analysis. By combining the study of the neurons from multiple lines of mouse brains, our data reveals the landscape of autophagy degradation in neurons and highlights selective sets of proteins from the ER and SV-related proteins as significant cargos of neuronal autophagy. We have identified Calumenin, an ER resident protein, as autophagy receptor and shown a novel function of autophagy in controlling PKA and neuronal activity through AKAP11. Our study provides a global view of autophagy degradation in human neurons and an insight into the mechanisms of neurological disorders linked to autophagy deficiency.

## Results

### Proteomic profiling of enriched cellular pathways in autophagy-deficient human induced neurons

Previous studies demonstrated that autophagy is constitutively active in neurons; basal autophagy constantly degrades cellular cargo to maintain neuronal homeostasis and protect neurons (Friedman et al., 2012; Komatsu et al., 2007a). To investigate the landscape of neuronal autophagy targets under basal conditions, we suspended autophagic flux in neurons, performed integrated proteomic profiling of accumulated proteins, and identified autophagy cargo, receptors, and targeted pathways (Figure 1A). To ensure the rigor of the analysis and isolate changes specific to autophagy inhibition, we elected to disrupt two essential autophagy genes, *ATG7* and *ATG14,* separately in human iNeurons as well as *Atg7* or *Atg14* in mouse brains in a neuron-specific manner. We also performed transcriptomic analysis by RNA sequencing (RNA seq) to discern the change of protein levels not caused by transcriptional alteration. Furthermore, we investigated the interactome of autophagy protein LC3 in the brains by including the green fluorescence protein-tagged LC3 (GFP-LC3) transgenic mice (Wang et al., 2006) in our study (Figure 1A).

**Figure 1.**
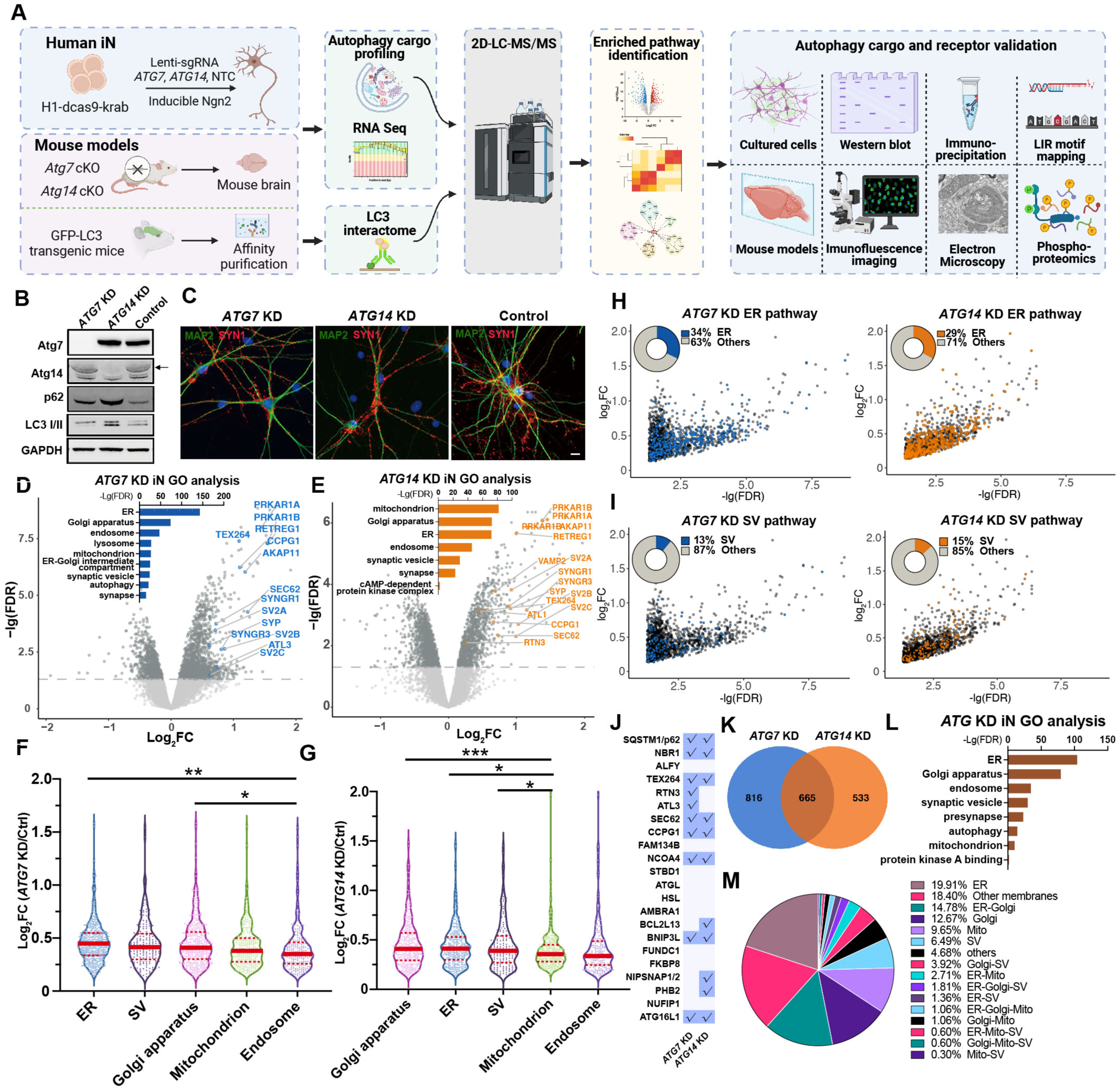
Proteomics analysis of human induced neurons (iNeurons) with *ATG7* or *ATG14* knock-down (KD) (A) An overview of the workflow for the integrated study of autophagy cargo in human iNeurons and mouse brains. (B) H1-dCas9-KRAB stem cells were used for the knock-down (KD) of *ATG7* or *ATG14* (arrow) and the block of autophagy was indicated by altered LC3I/II and p62 levels through immunoblot analysis. (C) Immunofluorescence images of *ATG7* KD and *ATG14* KD human iNeurons (6-week-old) co-stained with anti-MAP2 (green) and synapsin1 (SYN1, red) antibodies. Scale bar, 20 μm. (D) Volcano plot of the differentially expressed proteins (DEPs) detected by LC/LC-MS/MS in *ATG7* KD and control human iNeurons. Bar graph shows significant Gene Ontology (GO) annotations (FDR< 0.05, Log2FC> 0). FDR was calculated with the moderated *t*-test in the LIMMA package (R Studio). GO analyses were performed in g:Profiler (https://biit.cs.ut.ee/gprofiler/gost). (E) Volcano plot of the DEPs in *ATG14* KD and control human iNeurons. Bar graph displays the corresponding GO enrichments of the proteins (FDR< 0.05, Log2FC> 0). DEPs and GO analysis were performed as in (D). (F-G) Violin plots of DEPs enriched in ER, SV, Golgi apparatus, mitochondria, and endosome based on the GO analysis in *ATG7* KD and *ATG14* KD iNeurons, respectively. Each dot represents one protein. Solid red bars indicate the median Log2FC, and the dashed bars specify the 25th and 75th interquartile range. One-way ANOVA, *p<0.05, **p<0.01, ***p<0.001. (H) Plot of Log2FC and -Lg(FDR) for ER proteins of *ATG7* KD (blue, left) and *ATG14* KD (orange, right) (FDR< 0.05, Log2FC> 0) human iNeurons. Pie chart shows the percentage of ER DEPs and the rest from the mutant human iNeurons. (I) Plot of Log2FC and -Lg (FDR) for SV proteins of *ATG7* KD (blue, left) and *ATG14* KD (orange, right) (FDR< 0.05, Log2FC> 0) human iNeurons. Pie chart shows the percentage of SV proteins and the rest from the mutant human iNeurons. (J) A table of known autophagy receptors and those detected (check marks) in proteomic analysis of *ATG7* KD and *ATG14* KD human iNeurons (FDR< 0.05, Log2FC> 0). The list of autophagy receptors was manually collated through literature searches. (K) Venn diagram shows the overlap between the DEPs of *ATG7* KD (blue) and of *ATG14* KD (orange) iNeurons (FDR< 0.05, Log2FC> 0). (L) GO enrichment analysis of 665 DEPs shared between *ATG7* KD and *ATG14* KD human iNeurons (FDR< 0.05, Log2FC> 0). (M) Pie chart shows the percentage of proteins categorized into different organelles or pathways based on the dataset from (L).

Autophagy in human neurons is poorly characterized. We sought to investigate autophagy in human glutamatergic iNeurons, representing the most common neuron type in the CNS. We used our previously established approach to generate iNeurons by transiently expressing the transcription factor neurogenin 2 (NGN2) (Zhang et al., 2013) . Fusion of a catalytically inactive dCas9 to the KRAB repressor domain enables robust knockdown (KD) of endogenous genes in iNeurons (Tian et al., 2019). To repress *ATG7* or *ATG14* expression, we transduced human PSCs with a lentiviral construct expressing dCas9-KRAB and sgRNA targeting *ATG7* or *ATG14*. Reduction of ATG7 or ATG14 protein levels was robust in PSCs and iNeurons (Figure 1B, S1A and S1B). We observed no apparent effect of *ATG7* or *ATG14* KD on neuron growth or differentiation, as judged by neuron morphology or number at 6 weeks post-differentiation (PD) (Figure 1C). By this time, the iNeurons had obtained stable and robust electrophysiological properties and formed functional synapses (Zhang et al., 2013).

We next profiled the proteome of *ATG7* and *ATG14* KD iNeurons (6-week PD) by the multiplexed tandem mass tag (TMT)-based, two-dimensional liquid chromatography (LC/LC) and tandem mass spectrometry (MS/MS) analysis (Bai et al., 2017; Wang et al., 2020). We identified approximate 8,000 proteins across all the human iNeuron samples, demonstrating the high coverage of the proteome. Principal-component analysis (PCA) revealed reproducible replicate data, with the variance driven mainly by the difference in ATG gene expression. Similarly, clustering samples by protein expression similarity showed two main groups representing ATG KD and control iNeurons (Figure S1C and S1D). We identified 2372 and 2738 differentially expressed proteins (DEPs) in *ATG7* KD and *ATG14* KD iNeurons, respectively (FDR < 0.05). For the initial analysis of the accumulated proteins resulted from autophagy inhibition, we applied a relatively relaxed cutoff of a log2 fold change (Log2FC> 0; FDR< 0.05). Our quantitative analyses reveal 1481 and 1198 upregulated DEPs of proteins in *ATG7* KD and *ATG14* KD iNeurons, respectively, as shown in volcano plot analysis (Figures 1D and 1E). Gene ontology (GO) enrichment analysis of the upregulated DEPs highlighted cellular pathways associated with endoplasmic reticulum (ER), synaptic vesicle (SV), mitochondria, endosome, Golgi apparatus, and lysosome in both *ATG7* KD and *ATG14* KD iNeurons (Figure 1D and 1E). By plotting the fold increase of the upregulated DEPs, we obtained the mean values and inferred the abundance of the proteins for different organelles or pathways (Figure 1F and 1G). ER, SV, and Golgi DEPs noticeably displayed the greatest abundance in both *ATG7* and *ATG14* KD iNeurons relative to DEPs of mitochondria and endosome. Indeed, nearly one-third of the total upregulated DEPs are ER-related proteins in both *ATG7* and *ATG14* KD iNeurons (Figure 1H), while 13% and 15% of upregulated DEPs are SV proteins in *ATG7* KD and *ATG14* KD iNeurons, respectively (Figure 1I). Among the upregulated DEPs in *ATG7* KD and *ATG14* KD iNeurons, at least thirteen were known autophagy receptors as previously reported (Johansen and Lamark, 2020; Khaminets et al., 2016; Vargas et al., 2022), including five known ER-phagy receptors (Figure 1J). The identification of the previously described autophagy receptors validated feasibility and robustness of our approach.

To ask whether the increased levels of the proteins are due to transcriptional upregulation of their encoding genes, we performed RNA-seq analysis of *ATG7* KD iNeurons. Strikingly, *ATG7* was found as the only differentially expressed gene (DEG) (FDR<0.05) (Figure S1G). Furthermore, we found that 665 upregulated DEPs are shared between *ATG7* KD and *ATG14* KD iNeurons (Figure 1K), suggesting that they are unlikely *ATG7* or *ATG14* - specific but related to autophagy degradation. Consistently, these shared proteins were enriched in ER, Golgi apparatus, mitochondria, and SV-related pathways (Figure 1L and 1M). Noticeably ER proteins are top ranked as shown in both GO analysis (FDR) and the percentage of ER proteins in the upregulated DEPs (Figure 1L and 1M). Interestingly, we found the enrichment of “protein kinase A binding proteins” among the shared DEPs (Figure 1L).

Furthermore, we identified 822 and 1477 down-regulated proteins in *ATG7* KD and *ATG14* KD iNeurons, respectively (Log2FC <0, FDR < 0.05). 536 down-regulated DEPs are shared by the two mutant cells (Figure S1E). We found significant enrichment of the down-regulated pathways related to protein binding, synapse, and neuron development (Figure S1F).

### Impaired clearance of selective ER and SV proteins in autophagy-deficient human iNeurons

Our quantitative proteomics suggest that many ER proteins are significant autophagy cargo in human neurons, as indicated by high-ranking GO terms and abundance analysis in human iNeurons (Figure 1D, 1E). Multiple ER membrane proteins, including RTN3, ATL1, SEC62, and TEX264, function as receptors responsible for the selective degradation of the ER by autophagy (ER-phagy) (Chen et al., 2019; Chino et al., 2019; Fumagalli et al., 2016; Grumati et al., 2017). Immunoblot analysis confirmed that these ER-phagy receptors, along with ER resident proteins REEP5, Calnexin (CNX) and Calumenin (CALU), are increased or have a trend of increase in *ATG7* KD and *ATG14* KD iNeurons (Figures 2A, 2C and 2E). Immunofluorescence (IF) analysis showed a remarkable accumulation of CNX-labeled inclusions (arrows) particularly in the neurites of *ATG7* KD iNeurons (Figures 2G and 2H), suggesting an expansion of ER network. Using an ER-phagy reporter, which expresses an N-terminal ER signal sequence followed by tandem monomeric RFP and GFP sequences and the ER retention sequence KDEL (Chino et al., 2019), we found that the signal of the RFP was significantly reduced in *ATG7* KD iNeurons upon nutrient-withdrawal, indicating an impairment of ER-phagy (Figures S1H-S1J).

**Figure 2.**
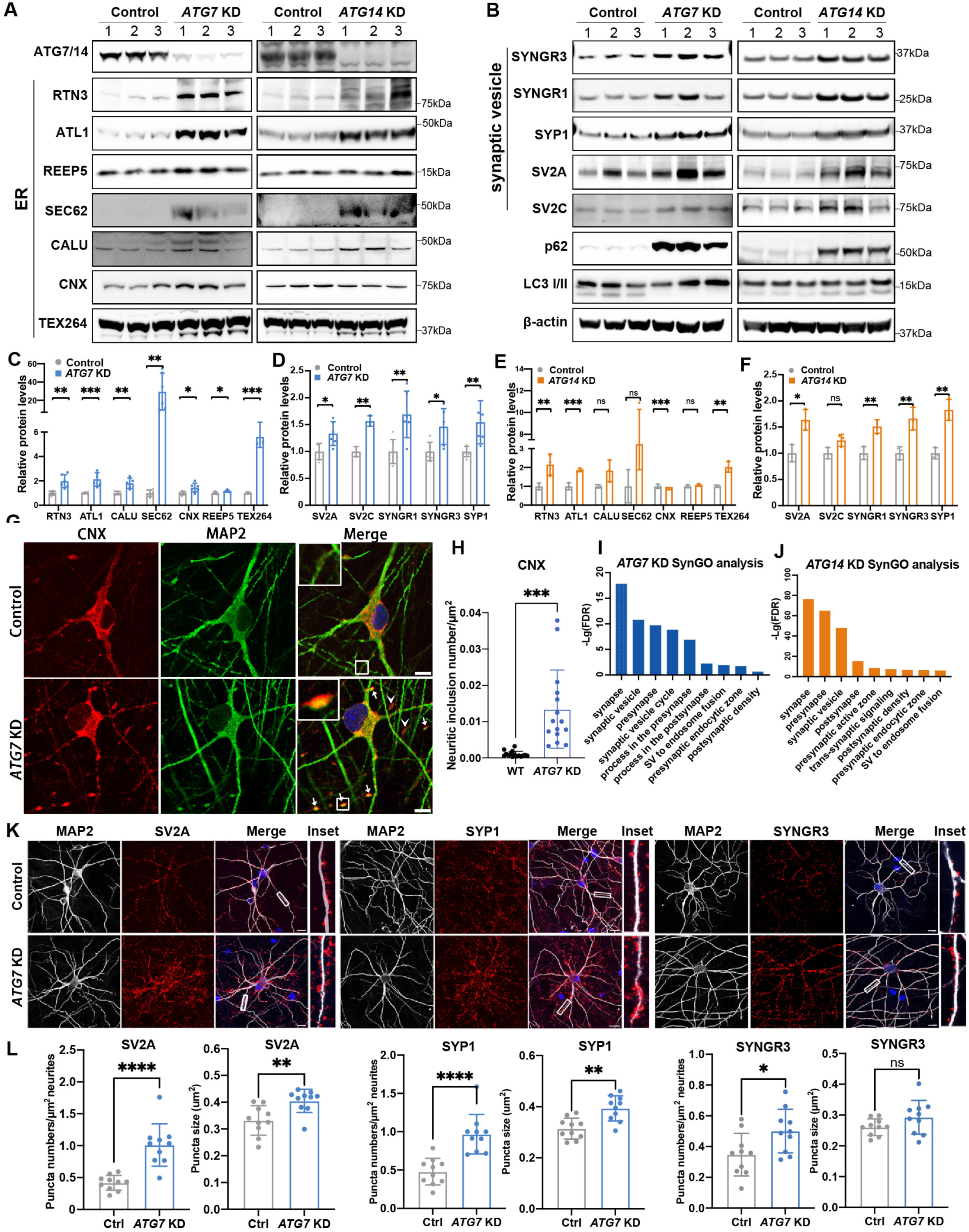
Validation of the DEPs of ER and SV related proteins in autophagy-deficient human iNeurons. (A) Immunoblot analysis of ER proteins as indicated in control, *ATG7* KD (left), and *ATG14* KD (right) human iNeurons. (B) Immunoblot analysis of SV proteins as indicated in control, *ATG7* KD (left), and *ATG14* KD (right) human iNeurons. (C-F) Quantification of the change of ER and SV proteins from (A) and (B). Relative protein levels were normalized to the loading control β actin. The controls’ means were set to 1. Unpaired t-test. Ns, no significance, *p<0.05, **p<0.01, ***p<0.001. (G) Immunofluorescence images of Calnexin (CNX, red) and MAP2 (green) staining in *ATG7* KD and control iNeurons. Insets, magnified images of the boxed region. (H) Quantification of CNX+ large inclusions per μ of total neuronal dendrite in *ATG7* KD iN and control iN. Unpaired t-test; Scale bar, 10μm . (n= 15) (I) Bar graph of top ranked GO terms generated by using SynGO from *ATG7* KD vs. control iN (FDR< 0.05, Log2FC> 0). (J) Bar graph of top ranked GO terms generated by using SynGO from *ATG14* KD vs. control iN (FDR< 0.05, Log2FC> 0). (K) Immunofluorescence images of *ATG7* KD human iNeurons stained with anti-SV2A, anti-synaptophysin1 (SYP1), and anti-synaptogyrin3 (SYNGR3), and anti-MAP2 antibodies. The highlighted areas were enlarged in inset. Scale bar, 10μm. (3 biological replicates, n=10 per batch) (L) Quantification of synaptic “puncta” density and size labeled with SV2A, SYP1, and SYNGR3 from (K). Unpaired t-test; Scale bar, 10 μm. (3 biological replicates, n= 10 per batch)

We next examined the accumulation of cargo candidates associated with SV. Significant increase of synaptogyrin1 (SYNGR1), synaptogyrin3 (SYNGR3), synaptophysin1 (SYP1), SV2A, and SV2C in autophagy-deficient iNeurons was confirmed by immunoblotting (Figures 2B, 2D and 2F). Further inspection of cargo candidates through SynGO analysis showed greater number of upregulated DEPs associated with presynapse than with postsynapse (Figures 2I and 2J), consistent with increased abundancy of autophagy cargo in presynaptic terminals (Goldsmith et al., 2022; Vijayan and Verstreken, 2017; Wang et al., 2006). IF staining revealed a significant increase in the size and density of SV2A-, SYP1-, and SYNGR3-associated puncta in *ATG7* KD iNeurons (Figures 2K and 2L), corroborating the clearance blockage of the above presynaptic proteins. We further confirmed the autophagy-lysosome degradation of SYNGR3, SV2A, SV2C, SYP1, and AKAP11 in iNeurons by bafilomycin A1 treatment, which causes an increase or trend of increase of these proteins (Figure S3B and S3C).

Collectively, our findings indicate that specific sets of ER and SV proteins require autophagy for the degradation in human neurons under basal conditions. ER-phagy is impaired in human *ATG7* KD iNeurons.

### Proteomic profiling of enriched cellular pathways in autophagy-deficient neurons from mouse brains

To validate our findings in human iNeurons, we sought to profile the targets of neuronal autophagy in vivo by using mouse brains. We selectively deleted *Atg7* or *Atg14* in CNS neurons using the combination of synapsin1 promoter-driven Cre recombinase mice (SynCre) and the *Atg7*f/f or *Atg14*f/f mice to generate conditional knock-out (cKO) mice. Immunoblot analysis of whole brain lysates from *Atg7*f/f-SynCre (*Atg7* cKO) and *Atg14*f/f-SynCre (*Atg14* cKO) mice confirmed the reduction of ATG7 and ATG14 protein levels, respectively, and an increase of autophagy substrate p62 and ubiquitinated protein levels (Figures S2A-S2D). Both autophagy-deficient mice exhibited a progressive reduction in body weight (Figures 3A and 3B) and viability (Figures 3C and 3D) over time, while *Atg14* cKO mice are more vulnerable than *Atg7* cKO mice. *Atg7* cKO and *Atg14* cKO brains (6-8 weeks old) were then subjected to the TMT-LC/LC-MS/MS analysis. We profiled 9,638 and 6,252 proteins from *Atg7* cKO and *Atg14* cKO mice, respectively. Most of the DEPs showed an increase in abundance (as opposed to a decrease) in the mutant brains (Figures 3E, 3F, S2E and S2F), suggesting a profound impact of autophagy deficiency on protein degradation. Indeed, 459 and 343 proteins were increased in *Atg7* cKO and *Atg14* cKO mouse brains compared to wild-type, respectively (*p* < 0.05, |Log2FC| > 2-fold standard deviation (2SD)) (Figures 3E and 3F). Like the findings in human iNeurons (Figure 1), the fraction of proteins whose abundance increased in the mutant mouse brains are enriched for GO terms, such as ER, mitochondria, Golgi apparatus, endosome, SV, autophagosomes, and cAMP-dependent kinase (PKA) (Figures 3E and 3F). Indeed, the upregulated DEPs of ER proteins account for 37% and 31% of the total upregulated DEPs from *Atg7* cKO and *Atg14* cKO brains, respectively (Figures 3I). We found that 102 upregulated DEPs are shared between *Atg7* cKO and *Atg14* cKO brains (Figure 3G), while ER components are the most significant in GO term analysis (Figure 3H). We detected increased abundance of SV proteins as well as cAMP-dependent protein kinase (PKA) pathways in both *Atg7* cKO and *Atg14* cKO mouse brains but to a lesser degree than ER or Golgi proteins (Figures 3E, 3F, 3I and 3J). Known ER-phagy receptors, such as TEX264, SEC62, and FAM134B, were found among upregulated DEPs in *Atg7* cKO or *Atg14* cKO mouse brains (Figure 3K). Furthermore, transcriptomic analysis using RNA-seq of *Atg7* cKO brains did not reveal an overlap between the upregulated DEGs and DEPs in *Atg7* cKO brains (Figure S2I and 3E). Instead, upregulated DEGs are activated glial marker genes, such as Gfap, C1qc, Clec7a, and Tyrobp (Figure S2I), potentially as responses to the dysfunctional or degenerating mutant neurons. Thus, the increase of protein abundance in *Atg7* cKO brains is unlikely due to the increase in transcriptional upregulation.

**Figure 3.**
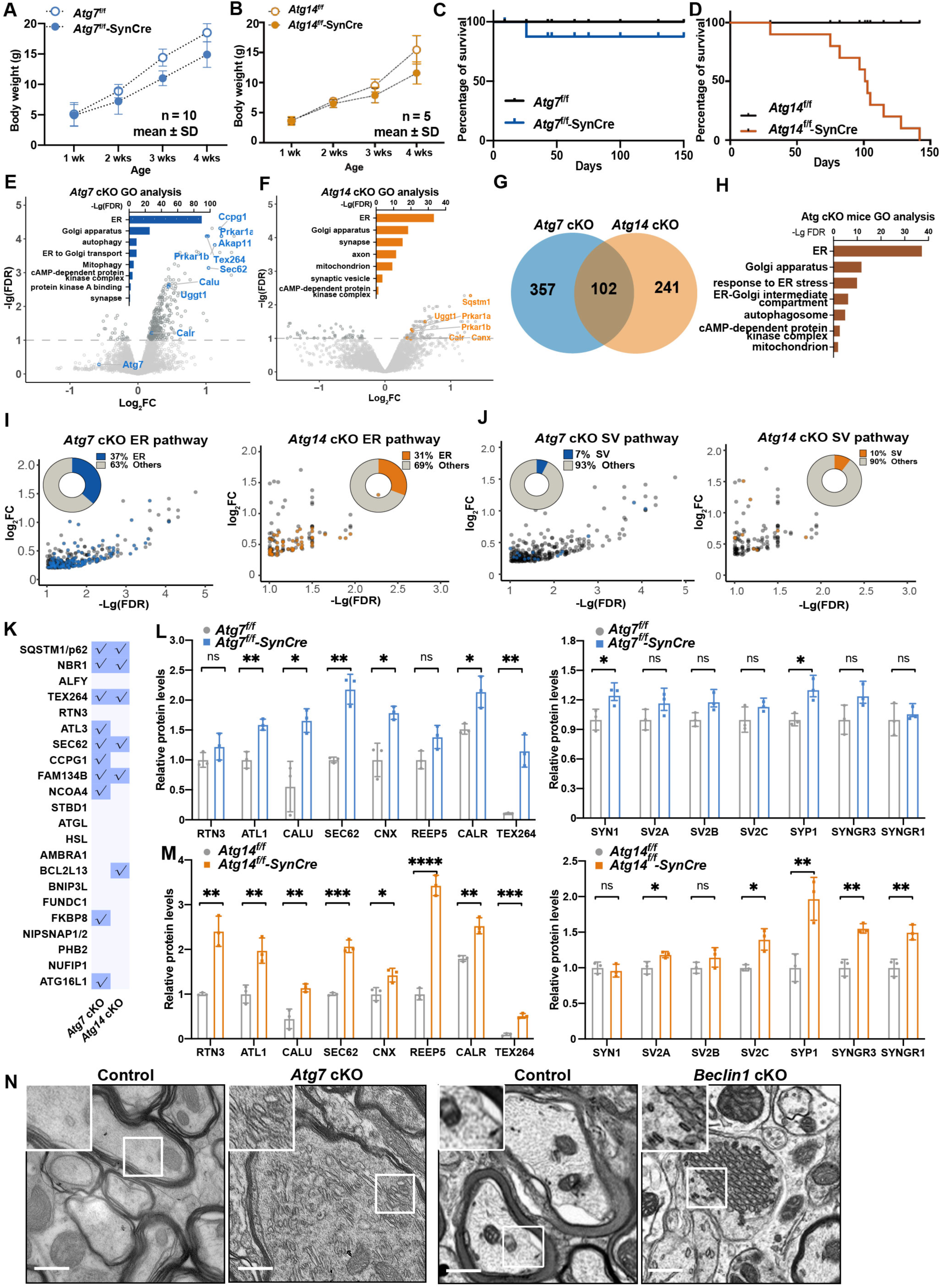
Proteomic analysis of proteins from mouse brains with neuron-specific KO of *Atg7* or *Atg14*. (A-B) Graph depicting body weight (grams) of *Atg7*f/f and *Atg7*f/f-SynCre mice (A), *Atg14*f/f and *Atg14*f/f-SynCre mice (B) at different ages. Data points represent mean +/-SD. (C-D) Kaplan Meier survival curve of *Atg7*f/f and *Atg7*f/f-Syn-Cre (C), *Atg14*f/f and *Atg14*f/f-Syn-Cre (D) mice at different ages. (E) Volcano plot of the DEPs from proteomic data of *Atg7*f/f and *Atg7*f/f-SynCre mouse brains. Inset bar graph shows the enriched GO terms associated with DEPs (*p* < 0.05, | Log2FC |>2SD, Log2FC> 0). (F) Volcano plot of the DEPs from proteomic data of *Atg14*f/f and *Atg14*f/f-Syn-Cre mouse brains. Inset bar graph shows the enriched GO terms among DEPs (*p* < 0.05, | Log2FC |>2SD, Log2FC> 0). (G) Venn diagram shows the overlap between the DEPs of *Atg7* cKO (blue) and of *Atg14* cKO (orange) mouse brains (*p* < 0.05, | Log2FC |>2SD, Log2FC> 0). (H) GO enrichment analysis of 102 DEPs shared between *Atg7* cKO (blue) and of *Atg14* cKO (orange) mouse brains (*p* < 0.05, | Log2FC |>2SD, Log2FC> 0). (I-J) Plot of Log2FC and -Lg (FDR) for ER (I) and SV (J) DEPs from *Atg7* cKO and *Atg14* cKO mouse brains, respectively. DEPs are indicated by blue (*Atg7* cKO) and orange dots (*Atg14* cKO). Inset circle plots show the percentage of ER DEPs vs. the rest of the DEPs. ER and SV proteins were defined by their GO annotations in gProfiler. (K) A table of the known autophagy receptors and those detected (check marks) in proteomic analysis of the *Atg7* cKO and *Atg14* cKO mouse brains (*p* < 0.05, | Log2FC |>2SD, Log2FC> 0. The list of autophagy receptors was manually collated through literature searches. (L-M) Quantification of the change of the indicated ER (left panels) and SV proteins (right panels) from *Atg7* cKO mouse brains vs. Control (L) and *Atg14* cKO mouse brains vs. Control (M). Relative protein levels were normalized to the loading control β actin. The controls’ means were set to 1. Unpaired *t*-test. NS, no significance, *p<0.05, **p<0.01, ***p<0.001, ****p<0.0001. (N) Electron microscopy (EM) images of Purkinje axons at the deep cerebellar nuclei (DCN) area from *Atg7*f/f and *Atg7*f/f-Pcp2-Cre mice (left panel) and *Beclin1*f/f and *Beclin1*f/f-Pcp2-Cre mice (right panel). Scale bar, 1 μm.

Consistent with human iNeuron results (Figure 1 and 2), immunoblot analysis demonstrated an increase or a strong tendency of increase in the abundance of ER proteins and ER-phagy receptors, such as RTN3, ATL1, SEC62, CNX, REEP5, and CALU, in *Atg7* cKO and *Atg14* cKO brains; and similar observations were found for SV proteins, including SV2A, SV2C, SYNGR1, SYNGR3, and SYP1 (Figures 3L, 3M, S2J and S2K). To determine if the ER protein accumulation is associated with ER structure alteration in mutant neurons, we performed electron microscopy (EM) analysis of *Atg7* cKO and *beclin 1* cKO brains. Like Atg14, Beclin 1 is a subunit of lipid kinase complex PI3K-III (VPS34) and is essential for autophagy (Zhong et al., 2009b). In both *Atg7* cKO and *beclin 1* cKO neurons, we found a remarkable expansion or stacking of tubular ER in the axons, corroborating with the increased levels of a specific set of ER proteins, such as tubular ER proteins RTN3 and REEP5, in mutant neurons (Figures 3N).

Taken together, the results from mouse brains largely validated the observations in human iNeurons and suggested that autophagy is required for the clearance of the proteins of multiple pathways and/or organelles in neurons, including ER, Golgi, mitochondria, SV, endosomes, lysosomes, and cAMP-PKA kinase complex. Our data indicates that ER proteins as well as tubular ER are the prominent targets of autophagy in neurons. Autophagy also targets a specific set of SV proteins in presynaptic terminals.

### Investigation of LC3-interactome in autophagy-deficient neurons

LC3 is the best-characterized mammalian Atg8 homolog. The lipidated LC3 is typically localized at autophagosomes and mediates cargo recruitment for degradation (Tanida et al., 2008). Green fluorescence protein (GFP)-tagged LC3 has been widely used to track autophagosomes in cultured cells and animal tissues (Bampton et al., 2005; Mizushima, 2009). GFP-LC3 transgenic mice were used to investigate LC3-binding proteins in mouse brains (Wang et al., 2006). Under basal conditions, however, GFP-LC3 is rarely detected in puncta or association with autophagosomes in the CNS neurons (Mizushima et al., 2004; Wang et al., 2006), possibly due to rapid cleavage of GFP-LC3 (Geng and Klionsky, 2017). To facilitate the identification of LC3-interacting cargos using anti-GFP affinity isolation, we inhibited neuronal autophagy in GFP-LC3 transgenic mice by crossing GFP-LC3 mice with the *Atg7*f/f-Syn-Cre mice (GFP-LC3; *Atg7*-cKO) and then performed GFP-pull-down from GFP-LC3; *Atg7*-cKO brains (Figure 4A). Immunoblot and silver staining confirmed the enrichment of proteins with GFP affinity purification, particularly in the GFP-LC3; *Atg7*-cKO mice (Figure S4A). Mass spectrometry analysis was performed to identify GFP-LC3 binding proteins remaining on the beads. The hierarchical clustering analysis confirmed similarity across biological replicates (Fig S4B). Next, we filtered out non-specific binding proteins by comparing GFP-LC3 and non-transgenic mice and focused on interactions that were altered in *Atg7-cKO*. Compared with *Atg7*-WT mice, *Atg7*-cKO mice had differential GFP-LC3 interaction in 600 proteins, while 73.7% were increased and 26.3% were decreased (Figure 4B).

**Figure 4.**
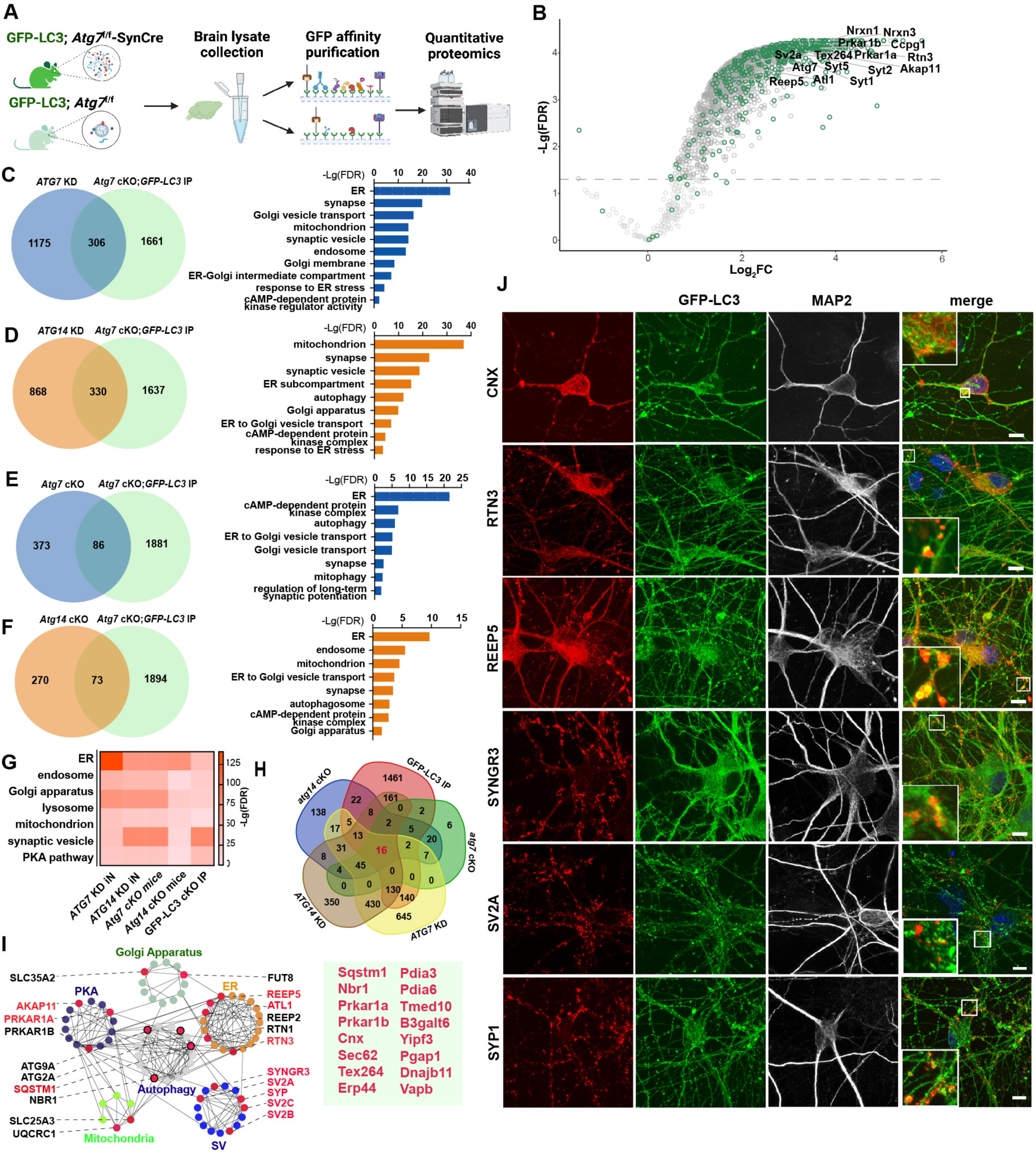
Proteomic analysis of GFP-LC3-interacting proteins from neuron-specific *Atg7* cKO mouse brains. (A) Schematic of investigation of LC3-interactome by combining genetic models (*GFP-LC3*; *Atg7*f/f or *Atg7*f/f-SynCre) and proteomics (TMT-LC/LC-MS/MS). (B) Volcano plot of proteomic results from DEPs between *Atg7*f/f-SynCre; GFP-LC3 IP vs. control IP (gray dots) and DEPs between *Atg7*f/f-SynCre; GFP-LC3 IP and *Atg7*f/f; GFP-LC3 IP (green dots). Dashed line indicates -Lg (FDR)= 1.3 (FDR= 0.05). (C) Venn diagram (left) of overlapping DEPs between *ATG7* KD human iN (blue) (FDR< 0.05, Log2FC> 0) and *Atg7*f/f-SynCre; GFP-LC3 IP (green) (FDR< 0.05, Log2FC> 0). Bar graph (right) displays the enriched GO annotations from the overlapping proteins. (D) Venn diagram (left) of overlapping DEPs between *ATG14* KD human iN (orange) (FDR< 0.05, Log2FC> 0) and *Atg7*f/f-SynCre; GFP-LC3 IP (green) (FDR< 0.05, Log2FC> 0). Bar graph (right) displays the enriched GO annotations from the overlapping proteins. (E) Venn diagram (left) of overlapping DEPs between *Atg7* cKO mouse brain (blue) (*p* < 0.05, | Log2FC |>2SD, Log2FC> 0) and *Atg7*f/f-SynCre; GFP-LC3 IP (green) (FDR< 0.05, Log2FC> 0). Bar graph (right) displays the enriched GO annotations from the overlapping proteins. (F) Venn diagram (left) of overlapping DEPs between *Atg14* cKO mouse brain (blue) (*p* < 0.05, | Log2FC |>2SD, Log2FC> 0) and *Atg7*f/f-SynCre; GFP-LC3 IP (green) (FDR< 0.05, Log2FC> 0). Bar graph (right) displays the enriched GO annotations from the overlapping proteins. (G) Heatmap for comparison of -Lg (FDR) of GO annotations enriched in *ATG7* KD iNeurons, *ATG14* KD iNeurons, *Atg7* cKO mice, *Atg14* cKO mice and *Atg7*f/f-SynCre; GFP-LC3 IP. (H) Venn diagram (top) of upregulated DEPs enriched in *ATG7* KD iNeurons, *ATG14* KD iNeurons, *Atg7* cKO mice, *Atg14* cKO mice and *Atg7*f/f-SynCre; GFP-LC3 IP. Numbers indicates protein numbers. 16 proteins shared by all the datasets were shown in the bottom panel. (I) Six examples of enriched PPI modules in upregulated DEPs shared by *ATG7* KD and *ATG*14 KD iNeurons with the proteins validated in this study highlighted in red. Each dot represents a protein, and the interaction is indicated by connected lines. (J) Immunofluorescence images of human iN producing GFP-LC3 with anti-ER proteins (CNX, RTN3, or REEP5) or anti-SV proteins (SYNGR3, SV2A, or SYP1). The cells were pretreated with chloroquine (100 μM) for 24 h before fixation. Insets, magnified images of the boxed regions. Scale bar, 10 μm.

We then performed pairwise comparisons between proteins with increased GFP-LC3 interaction in *Atg7* cKO brains and upregulated DEPs in *ATG7* KD iNeurons, *ATG14* KD iNeurons, *Atg7* cKO mouse brain or *Atg14* cKO mouse brains. The results revealed a considerable overlap of the increased protein levels (Figures 4C, 4D, 4E and 4F). GO enrichment analyses revealed that the molecular functions and cellular component of the overlapping proteins primarily comprised ER, Golgi trafficking, ER stress response, SV cycle, response to unfolded proteins, the autophagosome, and cAMP-dependent kinase (PKA) (Figures 4C, 4D, 4E and 4F). The reproducible enrichment of the PKA pathways as shown in the above multiple datasets is particularly intriguing, as we previously reported that autophagy selectively degrades PKA-RI subunits and regulates PKA activity through the AKAP11 receptor in tumor cells(Deng et al., 2021). Unsurprisingly, autophagy receptors (p62/SQSTM1, NBR1), ER-phagy receptors (TEX264, RTN3, SEC62, CCPG1), and mitophagy receptors (FUNDC1, NIPSNAP1/2) were detected in GFP-LC3 interactome validating our detection system (Figure S4D). By integrating GO enrichment pathways results from the above five datasets, we observed that the ER pathway is the most significant across most of the datasets, supporting the notion that ER is the most prominent target of autophagy in neurons (Figure 4G). We identified 45 proteins, which are shared by the four autophagy-deficient neuron datasets, and 16 shared by all five proteomic datasets (with LC3-interactome) of proteomics analysis (Figure 4H). These shared proteins include known autophagy cargo, such as SQSTM1, SEC62, NBR1, TEX264, PRKAR1A, and PRKAR1B. By performing protein-protein interaction (PPI) network analysis with selective upregulated DEPs shared by *ATG7* KD and *ATG14* KD iNeurons (665) (Figure 1K), we found specific and extensive interconnectivity among functional modules representing autophagy, ER, Golgi apparatus, SV, PKA pathway and mitochondrion (Figure 4I). Many components of the above modules were validated for their dependence on autophagy for clearance based on this study, including AKAP11, PRKAR1A, REEP5, ATL1, RTN3, SV2, SYNGR3, SYP (highlighted in red) (Figure 4I).

We next validated the interaction of LC3 with ER, SV, and PKA pathway proteins using the GFP-LC3 transgenic brains and iNeurons stably expressing GFP-LC3 (Figure S4C). Co-IP with GFP antibody confirmed interactions of LC3 with SV proteins (SV2A and SYNGR3) and AKAP11 in the mouse brain (Figure S4E). Moreover, we treated GFP-LC3 expressing iNeurons with chloroquine to block autophagy flux and performed IF analyses. Chloroquine apparently enhanced the number of GFP-LC3 puncta (indicating autophagosomes), which partially co-localize with the endogenous CNX, RTN3, and REEP5, SYP1, SV2A, and SYNGR3 (Figure 4J). These observations suggest that many LC3-binding proteins identified above are recruited to autophagosomes via LC3 for autophagy degradation in iNeurons.

### Identification of LIR-containing calumenin (CALU) as an ER resident autophagy receptor

Our integrated proteomics demonstrates a profound role of autophagy in targeting ER for degradation in human and mouse neurons under basal conditions. Six ER-phagy receptors have previously been reported in mammalian cells, including FAM134B (Khaminets et al., 2015), SEC62(Fumagalli et al., 2016), RTN3L (Grumati et al., 2017), CCPG1 (Smith et al., 2018), TEX264 (Chino et al., 2019), and ATL3 (Chen et al., 2019), which were all detected in our datasets including human iNeurons. Particularly TEX264 and Sec62 are significantly enriched in all 5 datasets in our study, suggesting a conserved ER-phagy function of TEX264 and Sec62 in human and mouse neurons.

Canonical autophagy receptors often harbor an LC3 interacting region (LIR), which binds to the LC3/GABARAP family of proteins (Kirkin et al., 2009). LIR motif consists of 4-6 amino acids, two of which are conserved (Birgisdottir et al., 2013; Jacomin et al., 2016). To identify novel autophagy receptors in neurons, we started from 341 proteins with significantly increased abundance (|Log2FC| >2SD, a stringent threshold) in human autophagy-deficient iNeurons. And then chose 180 ER-related proteins based on GO term pathway enrichment analysis to screen for putative LIR motifs (Jacomin et al., 2016), resulting in the identification of 45 proteins with transmembrane domains and at least one predicted LIR motif facing the cytosol. Several known ER-phagy receptors, including SEC62, CCPG1 and TEX264, were retrieved, validating our screening strategy. Interestingly, we identified calumenin (CALU) as a putative novel autophagy receptor and confirmed the increased CALU abundance in autophagy-deficient human iNeurons and mouse brains (Figures 2A, 2C, 2E, 3L, 3M and S2K).

CALU belongs to the CREC family of proteins containing EF-hand calcium-binding motifs and a signal peptide for endo/sarcoplasmic reticulum retention(Sahoo and Kim, 2010; Shen et al., 2019). To characterize CALU, we created *CALU* KO HEK293T and human iNeurons (Figures 5A, S5A and S5B). CALU staining with a specific antibody showed a reticular localization pattern and partially colocalizes with ER-resident protein CNX in iNeurons (Figures 5B and 5C). Furthermore, we transfected HEK293T cells with V5-tagged CALU (CALU-V5) expressing plasmid and found that CALU-V5 colocalizes with CNX and ER tubule marker RTN3 (Figure S5A). These results show that the CALU is an ER transmembrane protein in neurons. We verified the interaction of CALU and GFP-LC3B in HEK293T cells (Figure 5D) and in GFP-LC3B transgenic mouse brains through co-IP experiments (Figure 5E). Through IF analyses in iNeurons we found that CALU partially colocalizes with GFP-LC3B puncta, particularly in neurites, when the autophagy flux is blocked using chloroquine (Figure 5F). To test if the conserved LIR motif of CALU is required for LC3 binding (Figure 5G), we expressed hemagglutinin (HA)-tagged CALU-expressing plasmids with wild-type LIR (FIDL) or two mutated LIR (AIDA or AAAA) in HEK293T cells with the above plasmids. Co-IP showed that both LIR mutants completely abolish the interaction between CALU and LC3B, indicating that the FIDL sequence is a *bona fide* LIR in CALU required for LC3 binding (Figures 5G and 5H).

**Figure 5.**
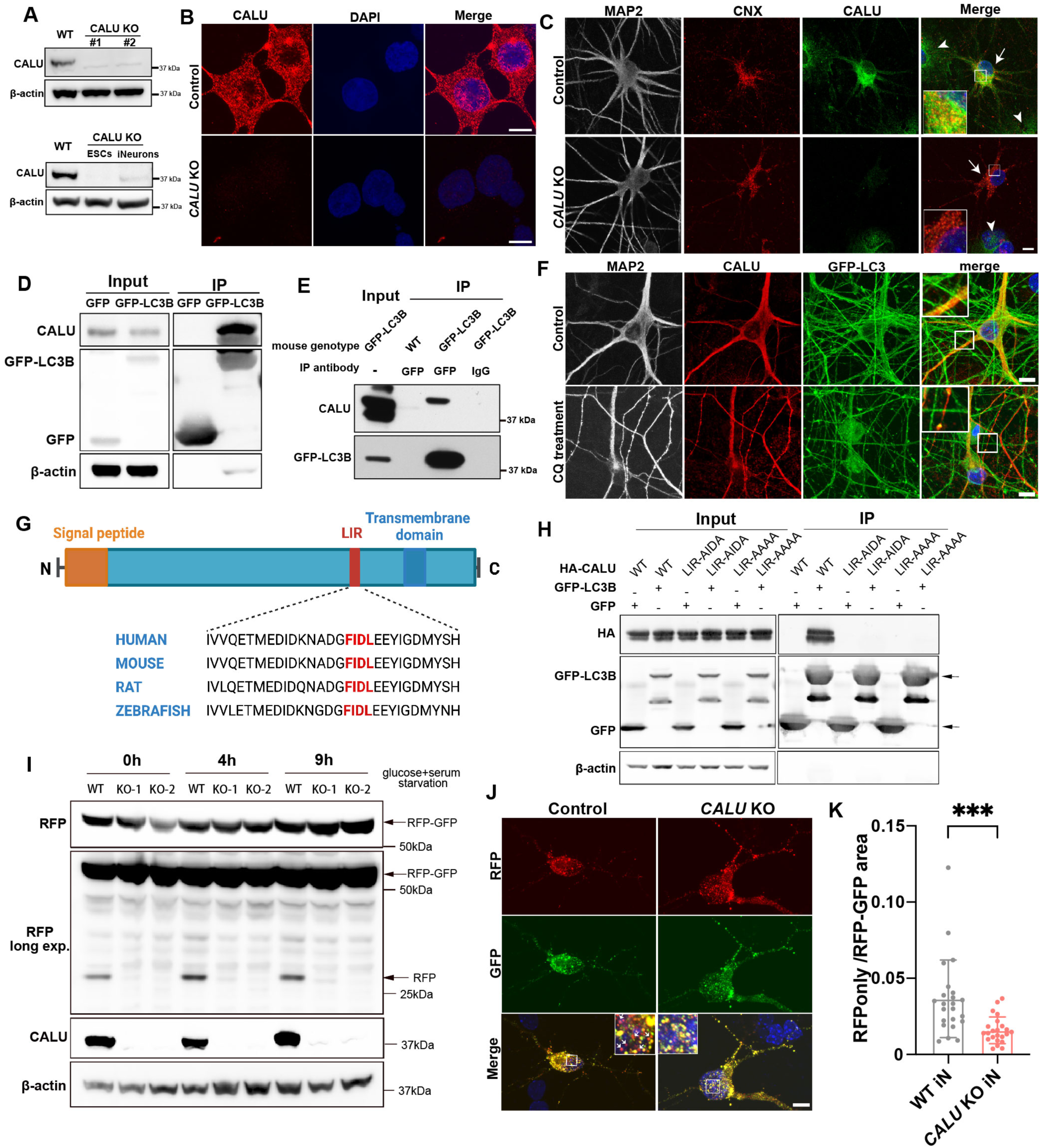
Characterization of LIR-containing calumenin (CALU) as a novel ER resident autophagy receptor. (A) Immunoblot analysis of CALU protein levels in two clones of *CALU* KO HEK293T cell line (top), *CALU* KO H1 stem cells, and the *CALU* KO stem cell-derived iNeurons (bottom). (B) Immunofluorescence imaging of *CALU* KO and WT HEK293T cells stained with an anti-CALU (red) antibody. Scale bar, 10 μm. (C) Immunofluorescence imaging of *CALU* KO and WT human iNeurons stained with anti-MAP2 (gray), anti-Calnexin (CNX, red), and anti-CALU (green) antibodies. White arrows indicate human iNeurons, arrowheads indicate mouse astrocytes (feeder). Insets, magnified images of the boxed regions. Scale bar, 10 μm. (D) Co-immunoprecipitation of endogenous CALU in HEK293T cells transfected with GFP-LC3B or GFP plasmids with anti-GFP antibody. (E) Co-immunoprecipitation of endogenous CALU with anti-GFP or control IgG antibodies in GFP-LC3 transgenic and WT mouse brains. (F) Immunofluorescence images of human iNeurons producing GFP-LC3 with anti-MAP2 (gray), anti-CALU (red), and anti-GFP (green) antibodies. The cells were pretreated with chloroquine (100 μM) for 24 h before fixation. Insets, magnified images of the boxed regions. Scale bar, 10 μm. (G) Schematic of CALU protein domain structure and the alignment of the LIR (LC3-interacting region) motif in CALU from different species. (H) Co-immunoprecipitation of CALU variants and GFP-LC3 proteins with indicated antibodies from HEK293T cells co-transfected with GFP or GFP-LC3B with HA-CALU-WT or HA-CALU-mutLIR (AIDA or AAAA) plasmids. Arrows indicate GFP and GFP-LC3 proteins. (I) Immunoblot with indicated antibodies of WT and two clones of *CALU* KO (#1 & #2) HEK293T cells stably expressing the ER-phagy reporter after nutrient starvation (lacking glucose and serum in medium) for 4 h and 9 h. (J) Immunofluorescence images of WT and *CALU* KO human iNeurons transfected with ER-phagy reporter plasmids after nutrient starvation for 48 h. Scale bars, 10 μm. (K) The RFP puncta only density was quantified. Data were collected from >21 cells for each cell type from three independent biological replicates. Un-paired *t*-test. *p < 0.05 (J).

We next evaluated the role of CALU in ER-phagy using *CALU* KO HEK293T cells and human iNeurons with the ER-phagy reporter as described above (Figure S3D). In the control cells, when ER-phagy was activated by starvation, the RFP fragment was released by lysosomal hydrolases, while the GFP signal was quenched within the acidic lysosomal environment (Figure S1H). In contrast, the signal of the RFP fragment was significantly reduced in both nutrient-replete and starvation conditions in *CALU* KO HEK293T cells (Figure 5I), indicating compromised ER-phagy activity. Similarly, by introducing the ER-phagy reporter in human iNeurons, we observed decreased RFP-only IF signals in *CALU* KO human iNeurons compared to control iNeurons (Figures 5J and 5K), suggesting impaired ER-phagy. Taken together, the data indicate that CALU is ER resident autophagy receptor.

### Autophagy degrades AKAP11 and RI proteins and regulates PKA and neuronal activity in human iNeurons

The integrated proteomics and LC3-interactome analysis suggest that the components of the cAMP-PKA pathway are significant targets of autophagy in human and mouse neurons (Figures 4G, 4H and 4I). Indeed, both PRKAR1A (RIα) and PRKAR1B (RIβ) of PKA subunits are significantly enriched in all autophagy-deficient iNeurons and mouse brains (Figure 4I). The PKA holoenzyme is a tetramer consisting of two regulatory subunits (R-subunits, RI and RII) and two catalytic subunits (PKAc). Each R subunit binds and inhibits a catalytic subunit at the resting state (Kim et al., 2007). cAMP binding to the R-subunits releases and activates PKAc (Day et al., 2011; Gold et al., 2006). We previously reported that AKAP11 is an autophagy receptor that mediates the selective degradation of the RI subunit (not RII) and activate PKA pathway in tumor cells (Deng et al., 2021). Here we also showed the interaction of LC3 and AKAP11 in neurons (Figure S4E).

Given the important functions of cAMP-PKA in neurons (Dubnau et al., 2003; Lee et al., 2021), we next explored the role of autophagy in the regulation of PKA-dependent neuronal activity. Immunoblotting revealed that AKAP11, RI (but not RII), and PKA Catalytic α subunit (PKA-Cα) protein levels were significantly increased in the autophagy-deficient iNeurons (Figure 6A), confirming the role of autophagy in regulating the turnover of PKA signaling components in neurons under the basal conditions. Blockage of autophagic flux by chloroquine leads to the accumulation of RI proteins colocalized with autophagosome markers GFP-LC3 and p62 (Figure 6B and 6C). We also knocked down *AKAP11* gene expression in iNeurons and found a significant increase of RI levels through both immunoblotting and IF analysis, demonstrating that RI requires AKAP11 for autophagy degradation in iNeurons (Figures S6E and S6F). To investigate whether autophagy regulates PKA-dependent neuronal activity, we analyzed forskolin-dependent phosphorylation of CREB (p-CREB) responses and expression of the immediate early response gene c-FOS in control and *ATG7* KD iNeurons. Immunoblot analysis revealed a reduction of forskolin-induced p-CREB in *ATG7* KD iNeurons (Figure 6D). Intriguingly, we found attenuated forskolin-dependent c-FOS induction, suggesting a decrease of neuronal activity in *ATG7* KD iNeurons (Figures 6D, 6E, and S6A). IF analyses showed the reduced staining intensity of p-CREB and c-FOS in the nuclei of *ATG7* KD iNeurons (Figures 6F and 6G). Studies of the primary cortical mouse neurons derived from *Atg7* cKO mice showed the similar results as in human iNeurons (Figure S6B). Altogether, our data suggest a previously unknown role of autophagy in regulating PKA-dependent neuronal activity in human and mouse neurons.

**Figure 6.**
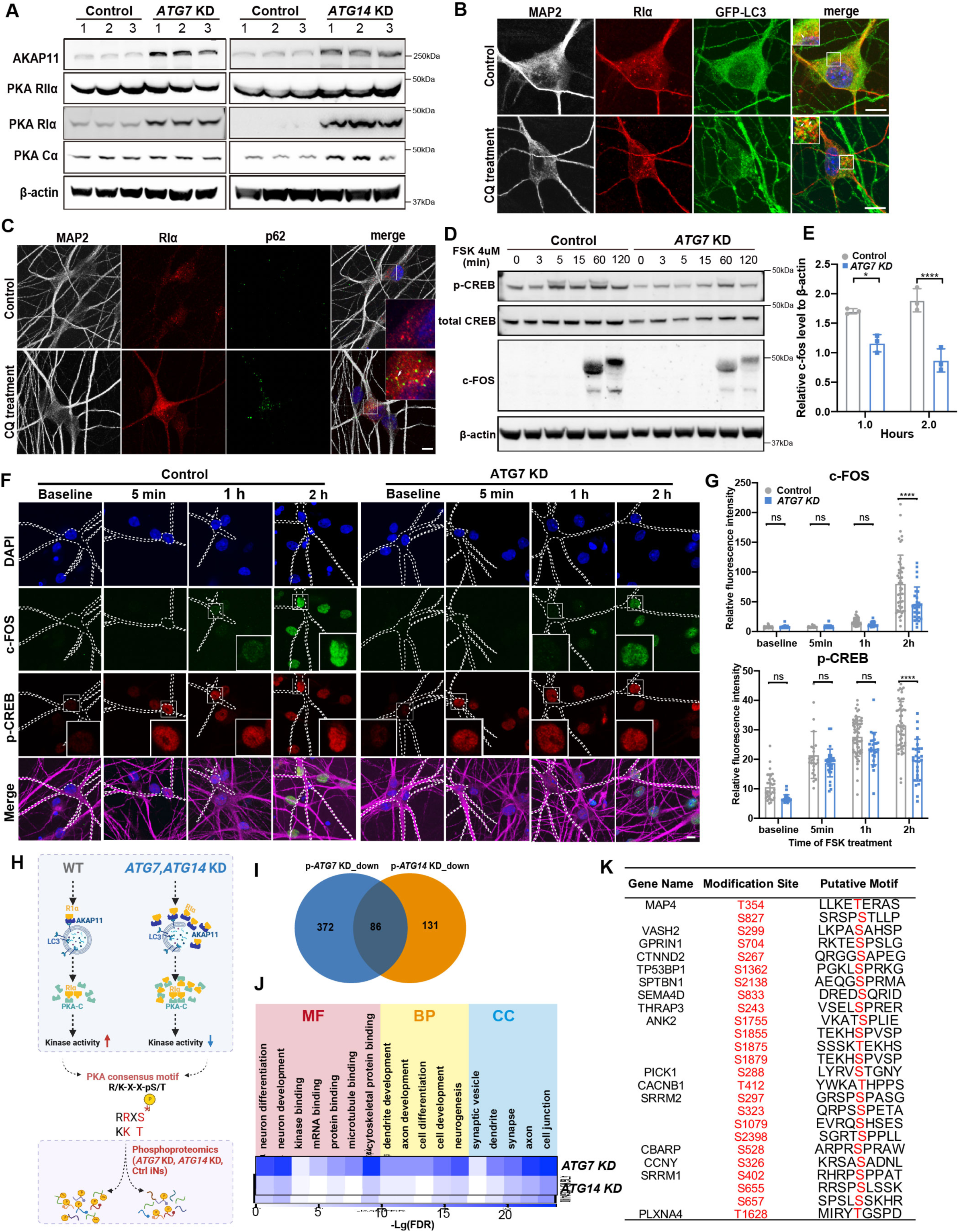
Characterization of AKAP11, PKA subunit levels and activity, and C-FOS level in human iNeurons with *ATG7* KD or *ATG14* KD. (A) Immunoblot analysis of AKAP11 and PKA subunits as indicated in control, *ATG7* KD (left) and *ATG14* KD (right) human iNeurons. Each lane represents an independent biological replicate. (B) Immunofluorescence imaging of PKA-RIα (red), GFP-LC3 (green) and MAP2 (gray) in human iNeurons expressing GFP-LC3. The cells were pretreated with chloroquine (100 μM) for 24 h to block autophagy. Insets, magnified images of the boxed regions. Scale bar, 10 μm. (C) Immunofluorescence images of PKA-RI Α (red), p62 (green), and MAP2 (gray) in control and *ATG7* KD human iNeurons. Insets, magnified images of the boxed regions. Scale bar, 10 μm. (D-E) Immunoblot analysis of CREB, p-CREB, c-FOS proteins in control and *ATG7* KD human iNeurons from i3N-derived iNeurons treated with Forskolin (4 μM) for 3, 5, 15, 60, 120minutes. Relative protein levels were normalized to β actin. Data were collected from three biological replicates. Un-paired *t*-test. *p < 0.05, ****p < 0.0001 (E). (F-G) Immunofluorescence images of c-FOS (green), p-CREB (red), MAP2 (magenta), and DAPI (blue) in control and *ATG7* KD human iNeurons treated with Forskolin (4 μM) for 0, 5, 60, 120minutes. The total signal intensity of c-FOS and p-CREB in MAP2-positive iNeurons were quantified. Scale bar, 10 μm. Data were collected from >25 cells for each cell type from three biological replicates. One-way ANOVA. Ns, no significance, ****p < 0.0001 (G). (H) Hypothetical changes of PKA subunit levels and activity and consequent alteration in the phosphorylation of PKA substrates in autophagy-deficient cells. (I) Venn diagram shows significantly downregulated phosphorylated peptides in *ATG7* KD and *ATG14* KD iNeurons (FDR < 0.1, Log2FC < 0). (J) Heatmap shows the -Lg (FDR) of enriched GO pathways for significantly downregulated phosphorylated peptides overlapped between *ATG7* KD and *ATG14* KD iNeurons using the dataset from (I). CC: Cellular Component; BP: Biological Process; MF: Molecular Function. (K) List of putative PKA substrates identified from the phosphoproteomics with PKA consensus motifs from overlapped dataset in (I).

PKA phosphorylates diverse downstream substrates to regulate critical functions in neurons (Zhong et al., 2009a). We reasoned that autophagy deficiency could lead to reduced phosphorylation of many PKA substrates (Figure 6H) due to the aberrant retention of the RI that sequesters the catalytic subunit. We then performed a phospho-proteomics analysis of *ATG7* KD and *ATG14* KD iNeurons. We identified 458 and 131 down-regulated phosphorylated protein sites in *ATG7* KD and *ATG14* KD iNeurons, respectively. There were 86 commonly phosphorylated sites between *ATG7* and *ATG14* KD iNeurons, of which many proteins carrying the phosphorylated sites were previously predicted as PKA substrates (Figure 6I). For example, phosphorylation of PKA substrates, including ADD1, ADD2, and NGFR(Dinkel et al., 2011; Higuchi et al., 2003; Matsuoka et al., 1996), were reduced in *ATG7* KD and *ATG14* KD iNeurons. The proteins with reduced phosphorylation in autophagy-deficient iNeurons showed strong enrichment in GO terms associated with neuronal functions such as neuronal cellular compartments, neurogenesis, and neuron development (Figure 6J). By screening for proteins with PKA phosphorylation consensus motif (R/K-X-X-pS/T)(Gesellchen et al., 2006) and associated with reduced phosphorylation in autophagy-deficient iNeurons, we identified 16 putative PKA substrates in the 86 shared down-regulated phospho-proteins (Figure 6K). Future studies will be needed to illustrate further their function in mediating neuronal activity.

### Loss of neuronal autophagy causes an aberrant increase of RI subunits and impairment of neuronal activity in mouse brains

Finally, we asked if autophagy-deficiency in neurons affects PKA subunit degradation and PKA, and PKAC Cα subunit levels are significantly increased in *Atg7* cKO and *Atg14* cKO mouse brains (Figures 7A and 7B). To explore the brain region specificity of the increased AKAP11 and RIα expression, we performed IF staining for AKAP11 and RI α in *Atg7* cKO and control brains. The overall fluorescence intensities of AKAP11 and RΙα staining were markedly increased across many brain regions of the mutant mice. For example, we found the accumulation of AKAP11 and RΙα-positive cytosolic aggregate-like structures in the hippocampus (CA3 and DG), cortex (layer IV), amygdaloid, and thalamic nucleus (Figures 7C-7H). Additionally, we found AKAP11 colocalized with p62 in the cytosolic aggregates in neurons (Figures 7E and 7F). IF staining of the mutant mouse brains revealed a reduced number of neurons labeled with c-FOS, indicating a decrease in neuronal activity at the basal level (Figures 7I and 7J). Together, our work suggests that the loss of autophagy disrupts PKA signaling in neurons and impairs neuronal activity *in vivo*.

**Figure 7.**
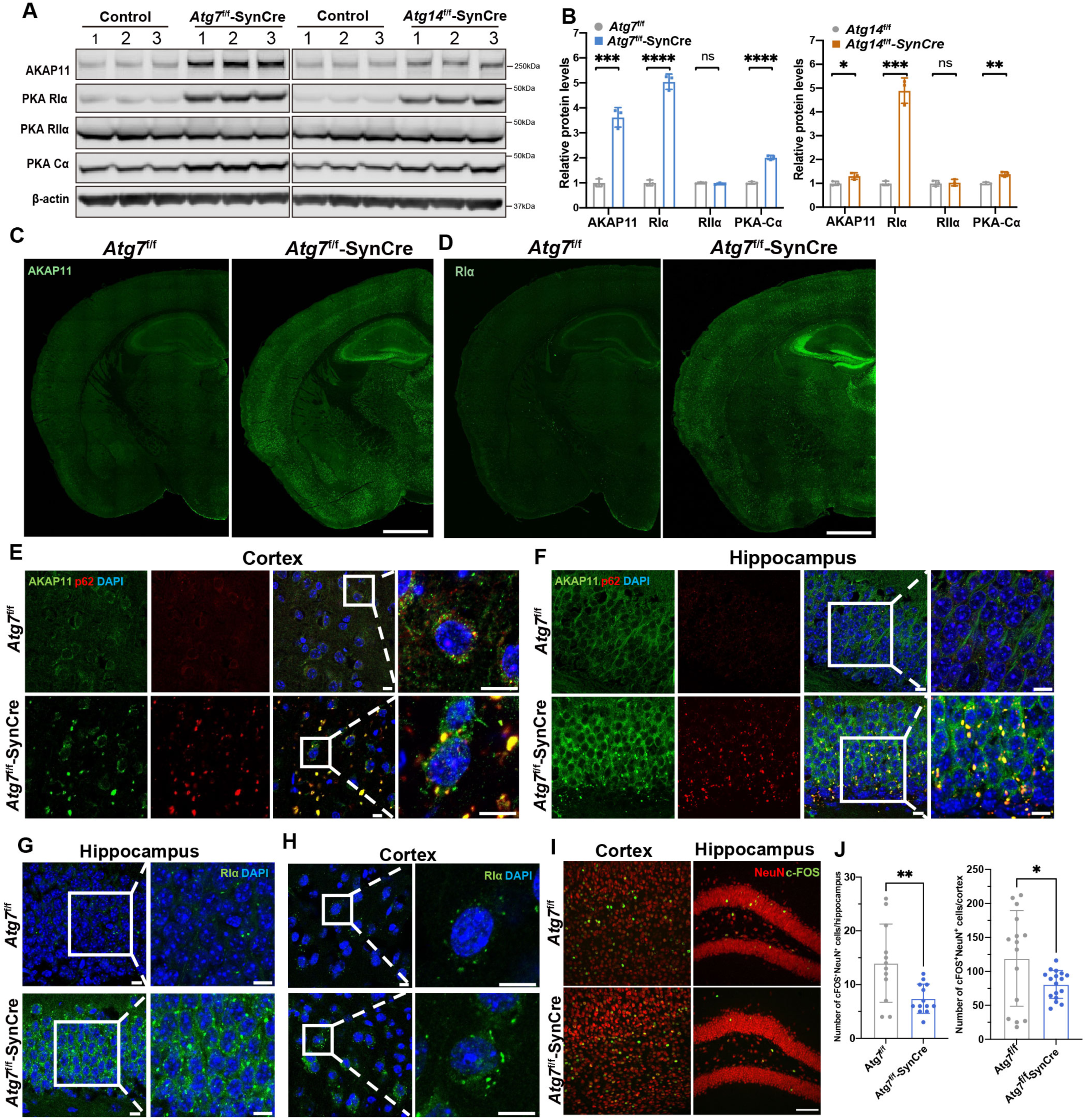
Examination of AKAP11, PKA RI subunit, and c-FOS levels in *Atg7* cKO mouse brains. (A) Immunoblot analysis of AKAP11 and PKA subunits as indicated in the whole brain lysates of *Atg7*f/f, *Atg7*f/f-Syn-Cre (left), *Atg14*f/f, and *Atg14*f/f-Syn-Cre (right) mice (2-month-old), n= 3. (B) Quantification of the change of proteins from (A). Relative protein levels were normalized to the loading control β actin. The controls’ means were set to 1. Unpaired *t*-test. *p < 0.05, **p < 0.01, ***p < 0.001, ****p < 0.0001. (C-D) Immunofluorescence images of *Atg7*f/f and *Atg7*f/f-SynCre mouse brain slices (2-month-old) stained with anti-AKAP11 (C) and anti-RIα antibodies. n=3, scale bars, 200μm. (E-F) Immunofluorescence images of *Atg7*f/f and *Atg7*f/f-SynCre mouse brain slices (2-month-old) co-stained with anti-AKAP11 (green) and anti-p62 antibodies (red) in the cortex (E) and hippocampus (F). n= 3, Scale bars, 10 μm. (G-H) Immunofluorescence images of *Atg7*f/f and *Atg7*f/f-SynCre mouse brain slices (2-month-old) stained with anti-RIα (green) antibody in the cortex (G) and hippocampus (H). n= 3, Scale bars, 10 μm. (I) Immunofluorescence images of *Atg7*f/f and *Atg7*f/f-SynCre mouse brain slices (2-month-old) co-stained with anti-c-FOS and anti-NeuN antibodies in the cortex and hippocampus (DG) at the basal level. n= 3 for each genotype, 3-5 slides for each brain region. Scale bars, 100 μm. (J) Quantification of c-FOS+NeuN+ neurons in (I). n= 3, 3-5 slides for each brain region.

## Discussion

Our study delineates the landscape of autophagy degradation in human and mouse neurons by applying integrated proteomic profiling and multidisciplinary experimental validation. We show that, under basal conditions, neuronal autophagy targets various cellular pathways and organelles, such as ER, mitochondria, Golgi, SV proteins, endosome, and cAMP-PKA pathway, for degradation. Our comprehensive analysis results in the identification of calumenin as a LIR-containing autophagy receptor. Furthermore, we show a novel function of neuronal autophagy in degrading PKA regulatory subunit RIα through the AKAP11 receptor, demonstrating a critical role of autophagy in regulating cAMP/PKA pathway and neuronal activity. Our study offers a valuable resource for understanding neuronal autophagy in human and insight into mechanisms of neurological disorders linked to autophagy deficiency.

Previous studies of potential autophagy cargo in neurons have provided useful information for the understanding of autophagy functions in neuron (Kuijpers et al., 2021; Ordureau et al., 2021; Zellner et al., 2021b). However, the discrepancy in the early findings led to inconsistent conclusions, which require further clarification and validation. Our study, differing from the previous studies, has significantly advanced the identification of autophagy targets and cargo in the following aspects: 1. unbiased, comprehensive approach with minimal perturbation of the systems; 2. larger proteome coverage and inclusion of multiple autophagy knockout models (human and mice), which enhance the rigor of the findings; 3. LC3-interactome analysis in the context of autophagy suppression to facilitate selective autophagy cargo capture; 4. interdisciplinary experimental validation of novel autophagy targets and receptors in neurons. Furthermore, the identification of many previously reported autophagy cargo and receptors in our datasets validates the robustness of our approach.

Among the many cellular pathways and organelles, ER emerges as the most significant target of neuronal autophagy based on our comprehensive analysis. The significance of ER-phagy in neurons has been suggested in previous studies by using *Atg5* cKO mice(Kuijpers et al., 2021) and human *ATG12*-deficient, NGN2-induced neuron(Ordureau et al., 2021), but contrasting the report of profiling AV with mouse brain(Goldsmith et al., 2022). The abundance and number of ER-related proteins are predominant among the GO enrichment pathways identified from autophagy-deficient human iNeurons and mouse brains as well as LC3 interactome analysis. Our results suggest that autophagy targets specific sets of ER proteins in human iNeuron neurites and degrades tubular ER in the axons of mouse brains. They demonstrate a profound function of neuronal autophagy in the maintenance of ER networks in the axon. Supporting the robustness of our approach for autophagy cargo identification, our results uncovered all six previously reported ER-phagy receptors, including TEX264, FAM134B, RTN3L, SEC62, CCPG1, and ATL3. Furthermore, our integrated approach promotes the identification of an ER-resident protein calumenin (CALU) as a novel autophagy receptor. CALU contains a transmembrane domain and six calcium-binding EF-hand motifs(Sahoo and Kim, 2010; Shen et al., 2019). This structural characteristic raises the question whether CALU plays a role in ER-phagy and calcium signaling of autophagic activity during autophagy. In addition, the role of CALU in autophagy regulation in neurons or axons remains to be clarified in the future.

Our analysis reveals that autophagy targets SV trafficking proteins for degradation in neurons. Our data for the first time suggests that autophagy targets a set of specific SV-related proteins, including synaptogyrin1 (SYNGR1), synaptogyrin3 (SYNGR3), synaptophysin1 (SYP1), SV2A, and SV2C in neurons. Binding of LC3 with SV2A and SYNGR3 also implicates the degradation through LC3-mediated autophagy. We note that SV2 proteins (including A, B, and C isoforms) contain the LIR in an anchor region of the cytosolic domain, which scores high in predicted interaction with LC3 protein. Future experiments should test the possibility that SV2 proteins act as autophagy receptors. Interestingly, although it was proposed that autophagy may recycle SV at presynaptic terminals(Vijayan and Verstreken, 2017), one study showed that disruption of autophagy has little impact on the number of SV, arguing against the idea that autophagy directly digest SV at the synapse(Kuijpers et al., 2021). Nonetheless, our observation provides an insight into the molecular mechanism whereby autophagy regulates presynaptic activity(Hernandez et al., 2012) by targeting selective SV proteins for degradation.

A striking finding of our integrated proteomics is the identification of cAMP/PKA pathway as a prominent target of autophagy in neurons even under basal conditions. Our results demonstrate that neuronal autophagy constantly degrades RI subunits of PKA holoenzyme and regulates PKA activity. Our previous studies found that AKAP11 acts as an autophagy receptor that mediates the selective degradation of PKA RΙα subunits in tumor cells(Deng et al., 2021). Our current study of AKAP11 KD iNeurons supports a role of AKAP11 as a selective autophagy receptor in mediating RΙα degradation in human neurons. We further show a remarkable AKAP11 and RΙα aggregation, which are colocalized with p62, in neurons at multiple brain regions, including the CA3 and DG regions of the hippocampus, layer IV of the cortex, amygdaloid, and thalamic nucleus from *Atg7* cKO mice. These observations suggest a particularly important role of autophagy in clearing RIα proteins and maintaining the homeostasis of PKA activity in these regions. Previous studies have shown that PKA is critical for mediating neuronal signaling pathways which is essential for learning and memory in mice and flies(Dubnau et al., 2003; Lee et al., 2021). Our findings thus help explain the mechanism whereby neuronal autophagy controls learning and memory(Lachance et al., 2019).

Finally, our study offers an insight into the molecular mechanism underlying multiple neurological disorders. A recent study reported 12 individuals from 5 independent families that carried biallelic, recessive variants in human *ATG7* associated with severe reduction or complete loss of ATG7 expression. These individuals suffered complex neurodevelopmental disorders (Collier et al., 2021). Our systemic study, particularly with human iNeurons carrying *ATG7* deletion, provides a comprehensive view for what cellular pathways and functions could be disrupted in the patients’ neurons carrying *ATG7* mutations. In Alzheimer’s disease (AD), dystrophic neurites and amyloid plaques are the important hallmarks of neuropathologies in the affected brains (Lee et al., 2022). Morphological and molecular analysis of AD brains and mouse models showed a sequential and distinct enhancement of staining of ATG9, RTN3, REEP5, RAB7, LC3, and LAMP1 proteins during dystrophic neurite and amyloid plaque development (Sharoar et al., 2019). The increased ATG9, RTN3 and REEP5 protein levels in AD neurites surrounds the early-stage amyloid plaques, suggesting that a block of ER-phagy occurs early and may contribute to the formation of dystrophic neurites and amyloid plaques in affected AD brains. Furthermore, rare AKAP11 truncating variants were recently identified as significant risk factors for both bipolar disorder and schizophrenia from exome sequencing (Palmer et al., 2022). Our observation raises a possibility that the pathogenic mechanism of the psychiatric disease involves disruption of cAMP-PKA activity and signaling in neurons.

In summary, our comprehensive characterization provides a global view of autophagy degradation in human and mouse neurons, offers a valuable resource of neuronal autophagy research, and shed light on the molecular mechanisms of several neurological disorders.

## AUTHOR CONTRIBUTIONS

Xiaoting Zhou conceived the study. You-Kyung Lee performed the sample preparation and immunoflueoscence staining with *Atg7* cKO mice brains. Xianting Li performed the proteomics sample preparation, immunoblot assay and GFP affinity purification using mouse brains. Henry Kim helped with the immunoflueoscence staining of human iNeurons, making schematics and data quantification. Carlos Sanchez-Priego generated the H1-dcas9-krab cell line. Xian Han, Haiyan Tan, Suiping Zhou performed the proteomics and phospho-proteomics analysis by MS. Yingxue Fu helped with the PPI protein network analysis. Kerry Purtell helped with the *Atg14* cKO mice breeding. Qian Wang performed the RNA sequencing data analysis, Gay Holstein helped with the EM analysis. Beisha Tang guided the experimental design. and Junming Peng guided the proteomics data analysis and data interpretation. Xiaoting Zhou, You-Kyung Lee, Nan Yang and Zhenyu Yue wrote the manuscript with input from all authors.

## ACKNOWLEDGMENTS

We are grateful to Ruiqi Hu for technical supports, Dr. Martin Kampmann for the kind donation of i3N cell line. We are also grateful to Dr. Insup Choi, Edward Wickstead, George Heaton and Marianna Liang in Yue’s laboratory for critical reading and discussion of the manuscript. The schematics were created using BioRender. This work was supported by the NIH grants R01NS060123, R01NS117590 and R21AG067570. X.Z was supported by China Scholarship Council and Central South University, Xiangya Hospital.

## Materials and Methods

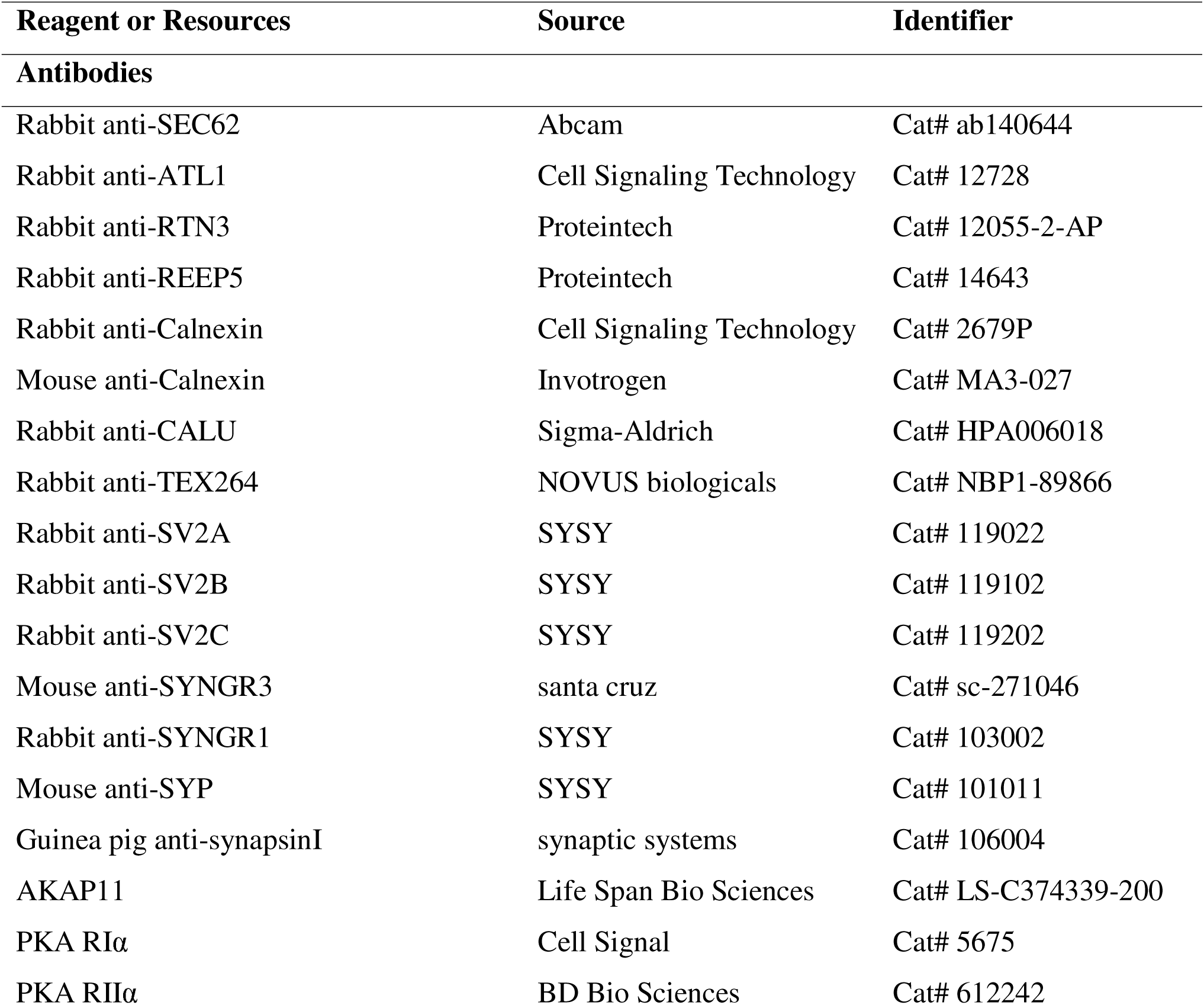
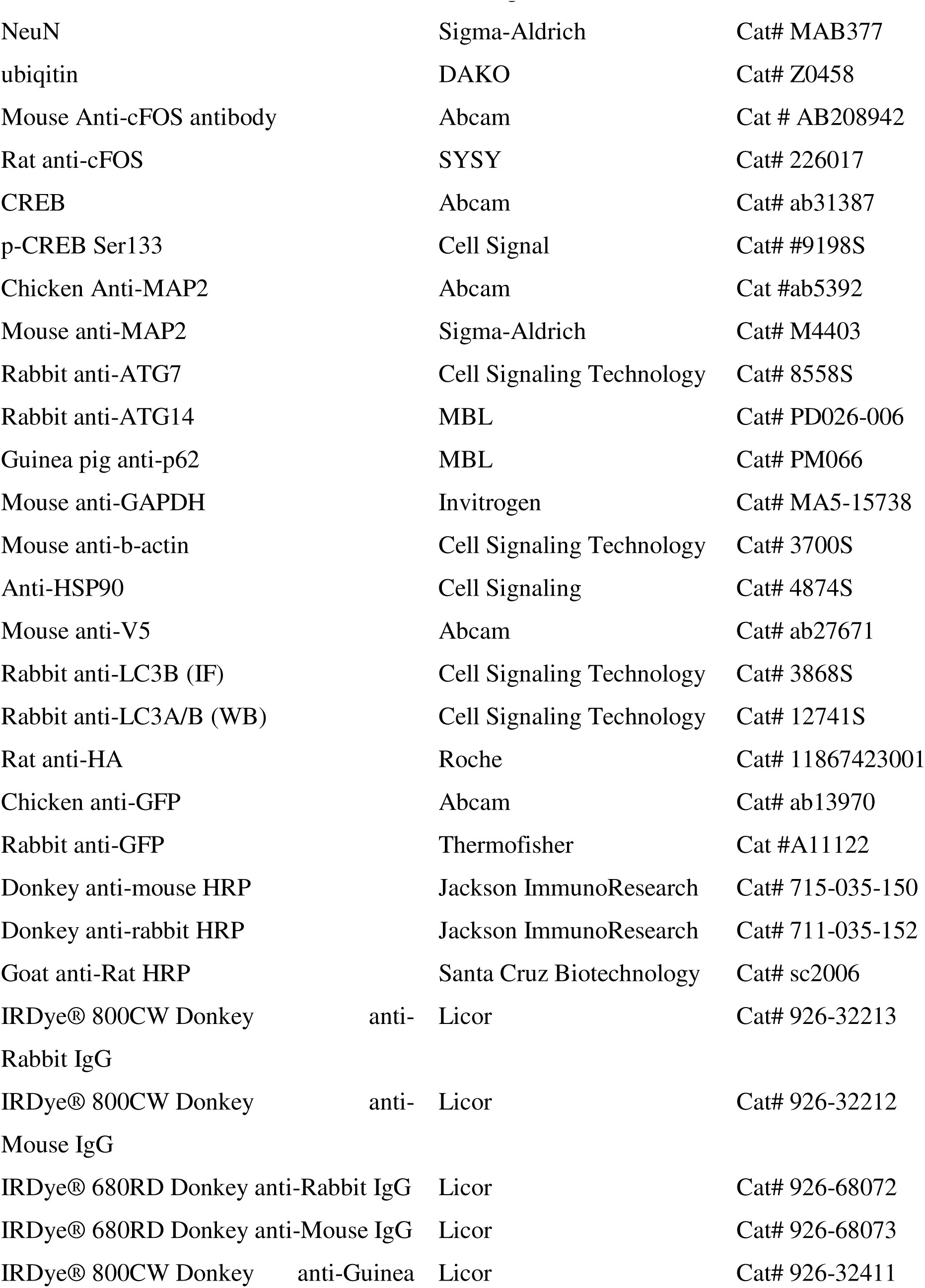
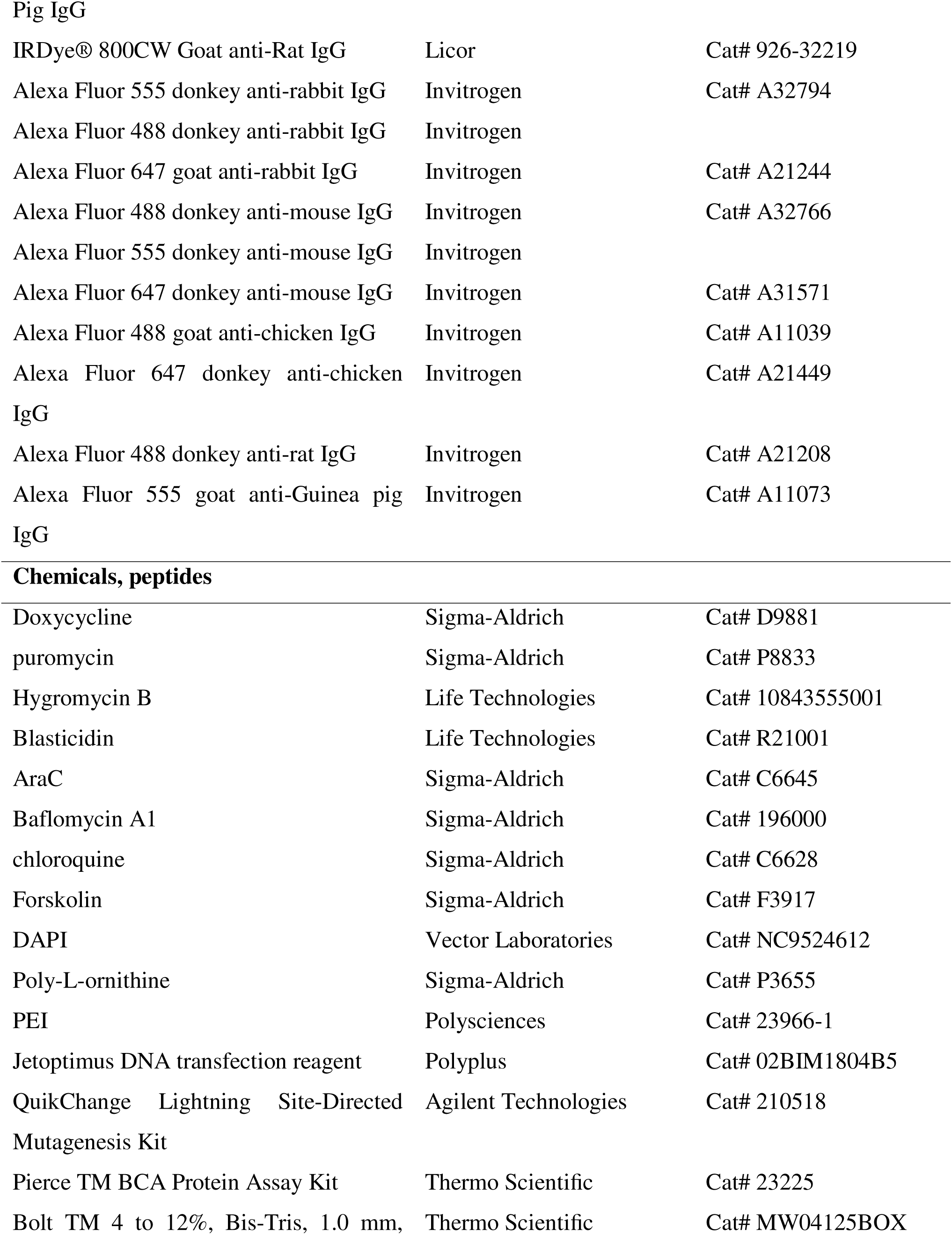
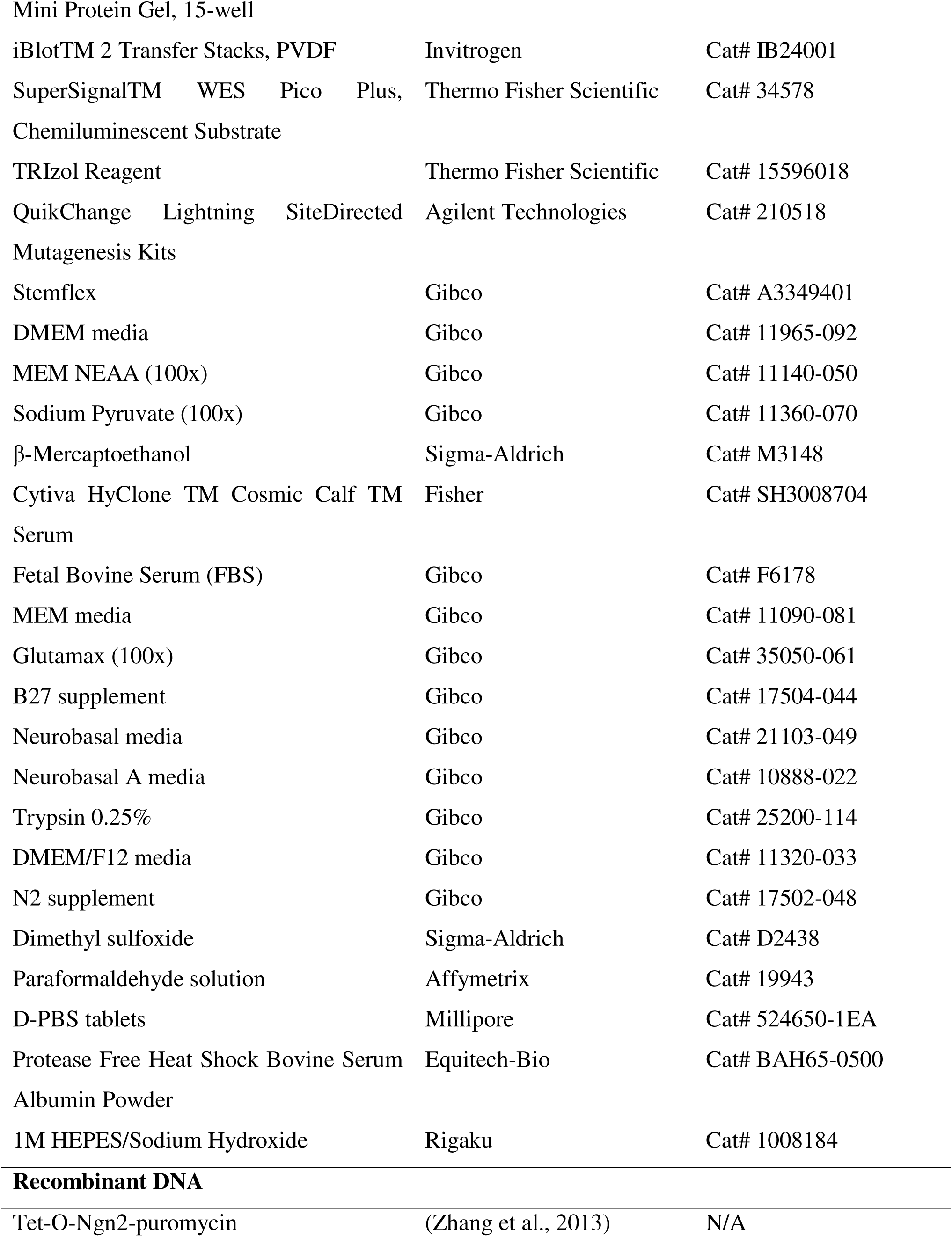
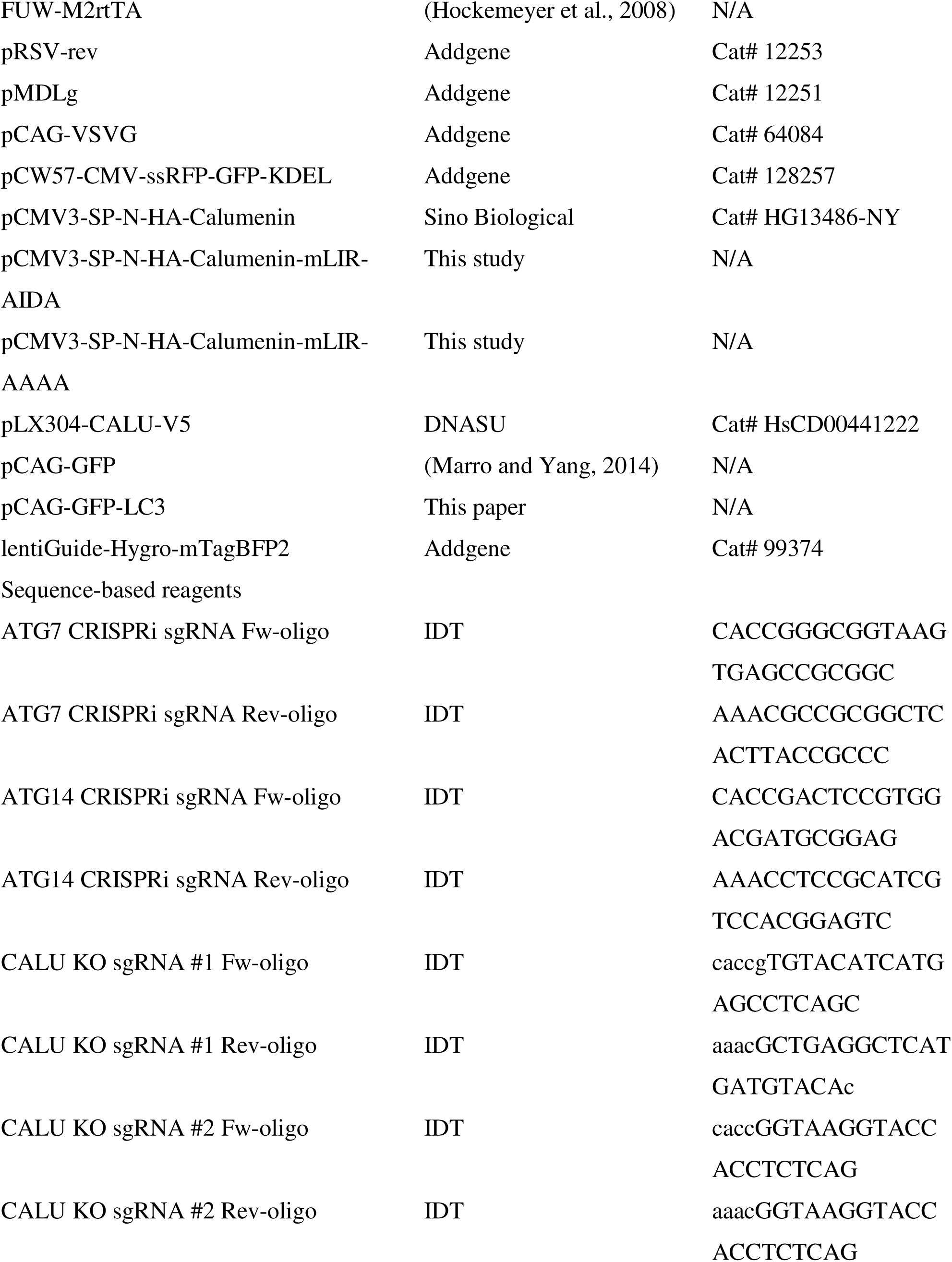
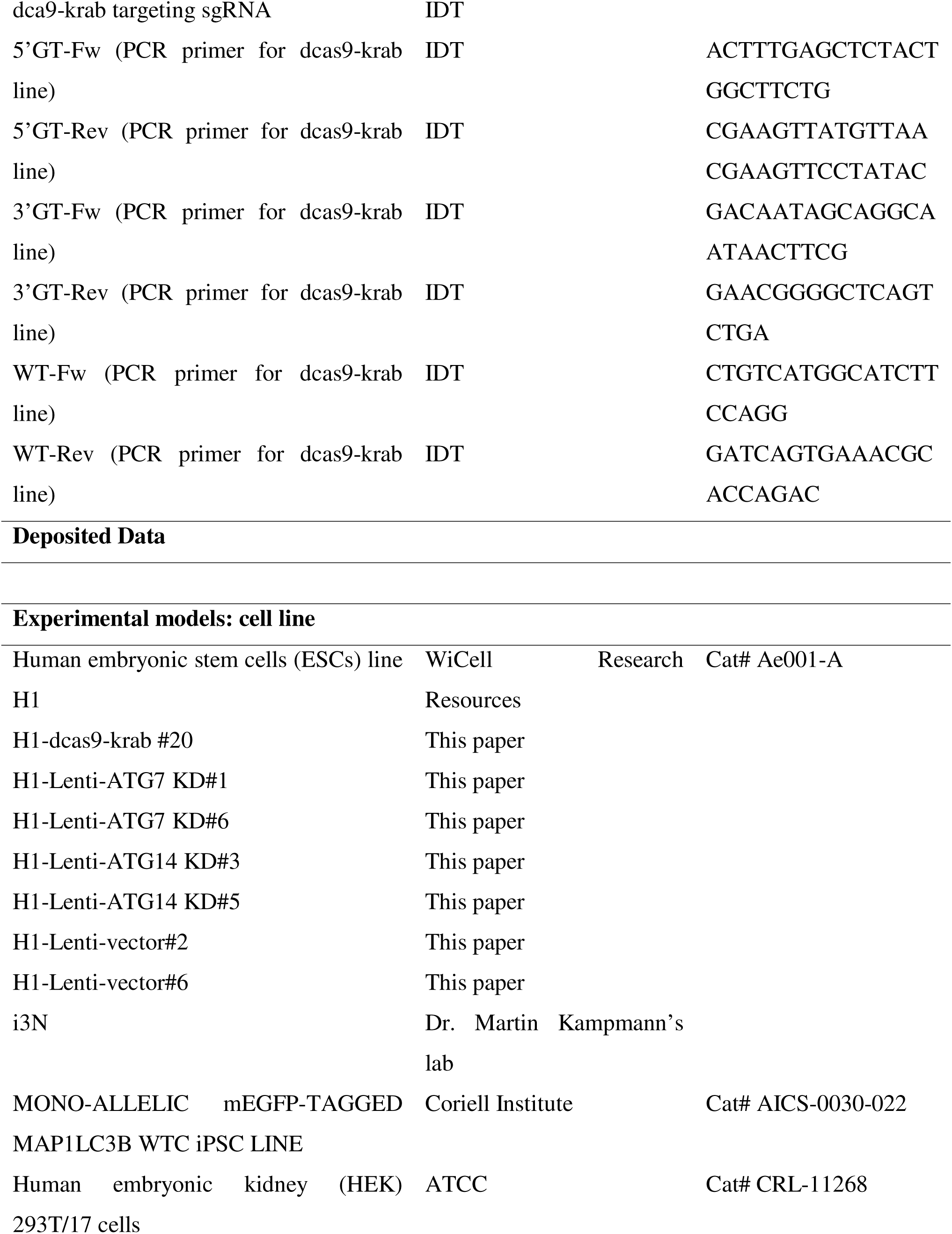
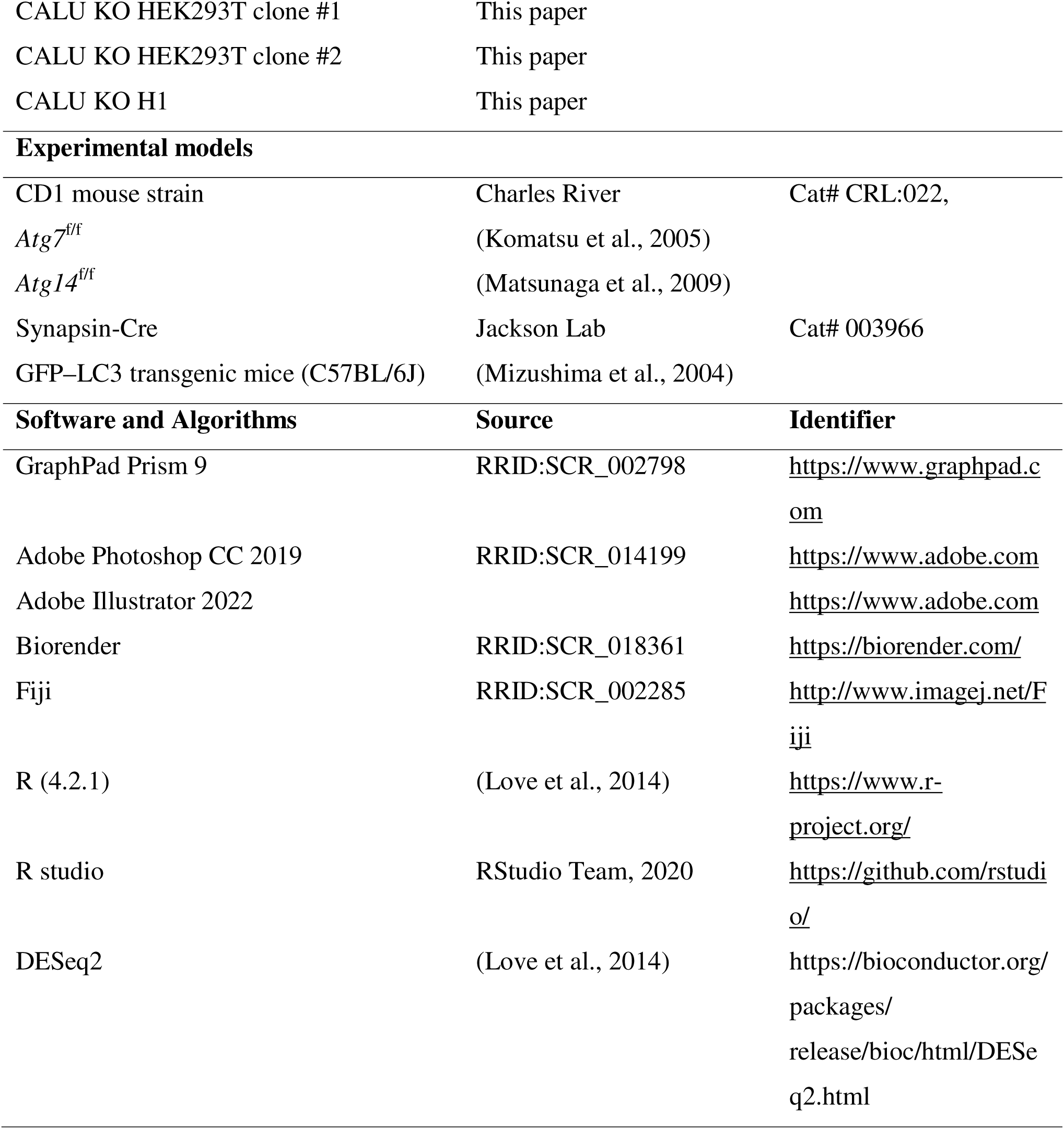

## RESOURCE AVAILABILITY

### Lead contact

Further information and requests for resources and reagents should be directed to and will be fulfilled by the lead contact, Zhenyu Yue, Nan Yang and Junmin Peng.

## EXPERIMENTAL MODELS AND SUBJECT DETAILS

### Animal models

All animal experiments were approved by the Icahn School of Medicine at Mount Sinai Institutional Animal Care and Use Committee and were conducted in compliance with the relevant ethical regulations. Mice were maintained in social cages on a 12h light/dark cycle with free access to food and water; animals were monitored daily for food and water intake. Wild-type CD1 mice were used to isolate primary cell cultures on postnatal day 3 (P3). Animals of both sexes were used in the analyses.

Floxed *Atg7* (*Atg7*f/f) and *Atg14* (*Atg14*f/f) mice were crossed with *Atg7*f/+-SynCre and *Atg14*f/+-SynCre mice to generate *Atg7*f/f-SynCre and *Atg14*f/f-SynCre mice. *GFP-LC3*; *Atg7*f/+-Syn-Cre mice were crossed with *Atg7*f/f mice to generate *GFP-LC3*; *Atg7*f/f-SynCre mice.

### Human cell lines

All human ESCs and PSCs were maintained on Geltrex-coated plates in feeder-free Stemflex medium and a 5% CO_2_ environment at 37□.

## METHODS DETAILS

### Cell culture

All human ESCs and PSCs were maintained on Geltrex-coated plates in feeder-free Stemflex medium and a 5% CO_2_ environment at 37□. Cells were passaged using 0.5 mM EDTA/PBS in Stemflex supplemented with 2 mM Thiazovivin or 50nM Chroman1. Thiazovivin/Chroman1 was removed from the medium on the following day. Research performed on samples of human origin were conducted according to protocols approved by the Institutional Review Boards of Icahn School of Medicine at Mount Sinai. H1 (WA01) ES cells were obtained from WiCell Research Resources; The inducible i3N iPS cell line was kindly provided by Dr. Martin Kampmann’s lab.

HEK293T cells were maintained in MEF medium (Dulbecco’s modified Eagle’s medium (Gibco, 11965-092) supplemented with 10% Cosmic Calf Serum (HyClone, SH30087.03), 1× NEAA (Gibco, 11140-050), 1×Sodium Pyruvate (Gibco, 11360-070), 0.008% β-mercaptoethanol).

Mouse glial cultures were generated from cortical hemispheres at postnatal day 3 (P3). The cortices from 3 pups were incubated in 5 mL of 20 Units/mL Papain, 0.5 mM EDTA, and 1 mM CaCl2 in HBSS for 15 minutes. After incubation, the tissues were manually dissociated by forceful trituration. Cells were filtered using a 70 μ cell strainer. The resulting cells were grown in MEF medium at 37°C with 5% CO_2_.

### sgRNA design and cloning to knock down *ATG7* and *ATG14* using CRISPR inhibition

4 small guide RNA (sgRNA) candidates for *ATG7* and 4 sgRNA candidates for *ATG14* were designed using the Broad Institute CRISpick web-based tool (https://portals.broadinstitute.org/gppx/crispick/public). sgRNAs were cloned into the Lenti-U6-dcas9-krab-Puro (Addgene #71236) plasmid using the Golden Gate DNA Assembly kit (NEB, E1602S). sgRNAs sequences were confirmed by Sanger sequencing. WT human H1 stem cells were transduced with lentiviruses overexpressing sgRNA-dcas9-krab-puro and puromycin (1ug/ml) was applied for at least 3 days to enrich puromycin resistant cells. Western blotting was performed to confirm the efficiency of *ATG7* and *ATG14* knock down. *ATG7* sgRNA#3 and *ATG14* sgRNA#3 were confirmed to have the highest knock down efficiency.

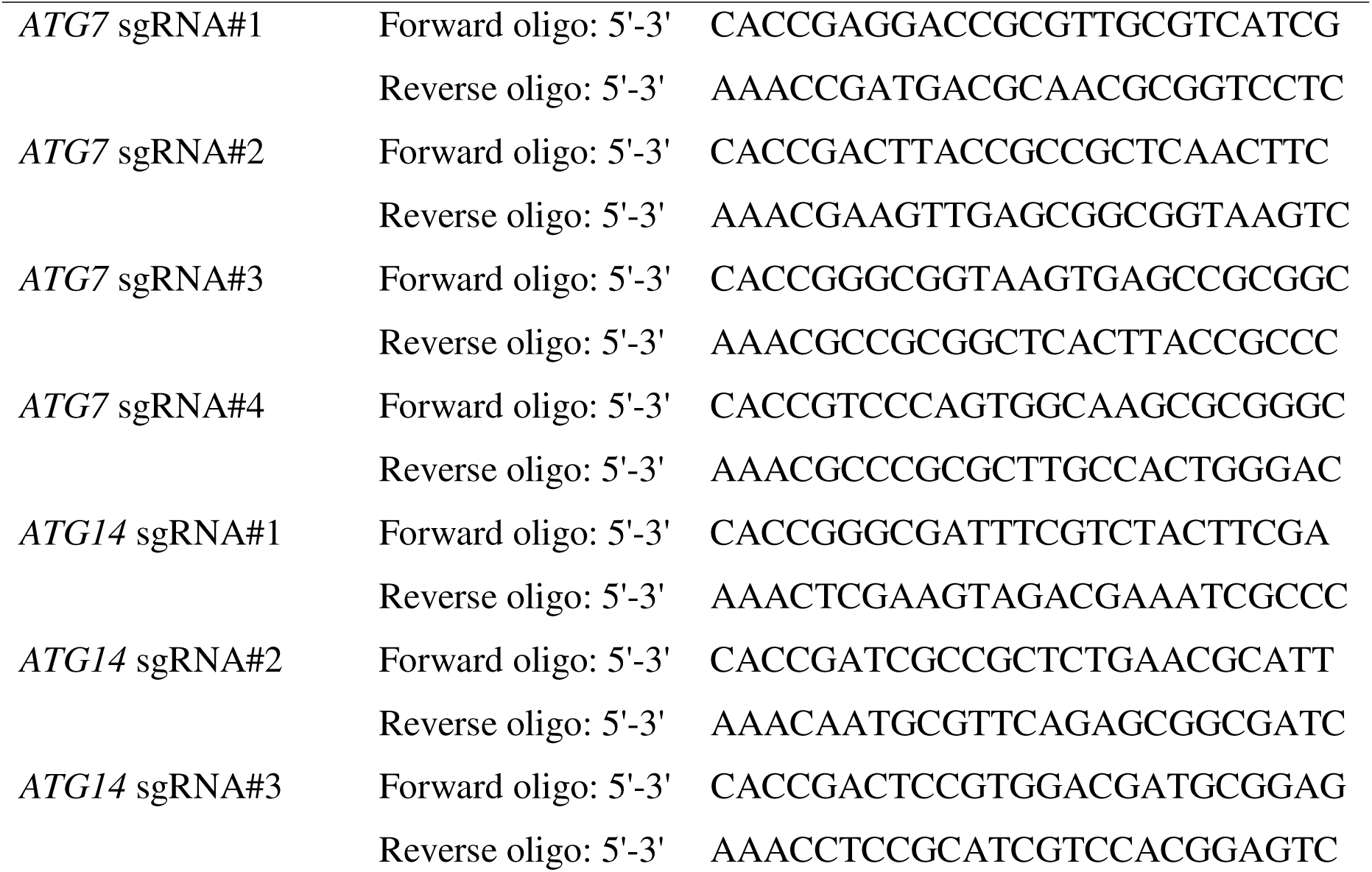
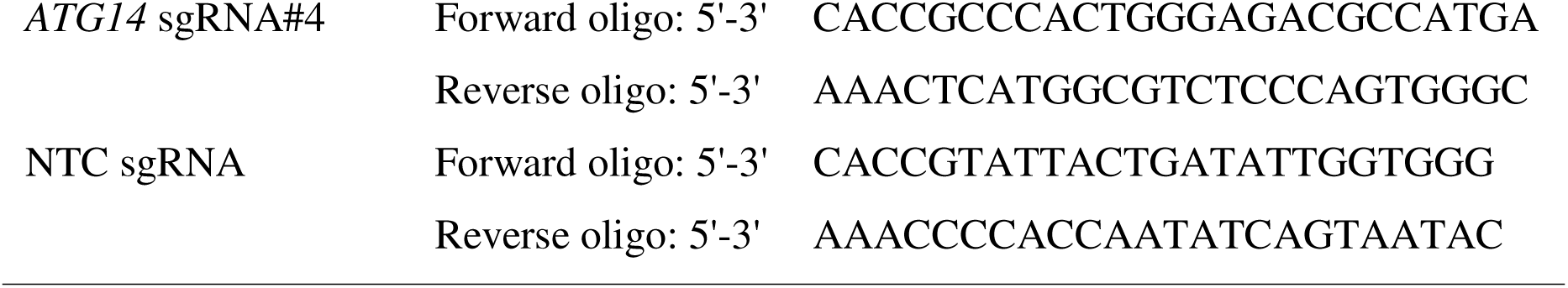

### Plasmid cloning

HA-*CALU*-mutLIR was constructed using a Quick-Change Lightning Site-Directed Mutagenesis Kit.

### Cell line generation

#### H1-Lenti-*ATG*7/14 KD cell line

Human H1 stem cells were dissociated using 0.5 mM EDTA/PBS and plated in Stemflex supplemented with 50 nM Chroman1. Lentiviruses (Addgene #71236) expressing *ATG7* sgRNA #3, *ATG14* sgRNA #3, or an empty vector was used to transduce H1 cells during passaging. The medium was changed to Stemflex supplemented with puromycin (1μg1ug/ml) for 3 consecutive days. Puromycin resistant cells were dissociated using Accutase for 15 minutes and seed at a density of 2,000 cells per 10 cm dish. Individual clones were harvested and expanded. Protein expression was confirmed by immunoblotting assay. *ATG7* KD clone #1 and #6, *ATG14* KD clones #3 and #5, and empty vector clones #2 and # 6 were confirmed by western blotting and used for subsequent experiments.

#### H1-dcas9-krab cell line generation

A FRT-EMCV-IRES-Neo-FRT-EF1a-dcas9-HA-krab transgene was subcloned into a pSIN donor vector containing an AAVS1 homology arm using CRISPR Cas9. Genomic DNA from Neomycin-selected cells were expanded, purified, and genotyped by three PCR reactions.

#### CALU KO cell line generation

The single guide RNA (sgRNA) to generate *CALU* KO H1 cells were synthesized (Integrated DNA Technologies) and cloned into lentiCRISPR version 2 plasmid. Human H1 ES cells were transfected with sgRNA plasmid pairs using Lipofectamine Stem Reagents (Invitrogen, Lot-01250336). The medium was changed to Stemflex supplemented with puromycin (1ug/ml) for 2 consecutive days. Puromycin resistant cells were dissociated using Accutase for 15 minutes and seeded at a density of 2,000 cells per 10 cm dish. Individual clones were harvested and expanded.

PCR products from each clone were sequenced and protein expression was confirmed by immunoblotting assay. Similarly, for the CALU KO HEK293T cell line generation, HEK293T cell were seeded at 30,000 cells/cm2 and transfected with the same pair of LentiCRISPR-v2 plasmids overexpressing sgRNAs using PEI 24 h after transfection. The medium was changed to MEF supplemented with puromycin (1ug/ml) for 3 consecutive days. Puromycin resistant cells were dissociated using 0.25% Trypsin for 5 minutes and seed at a density of 2,000 cells per 10 cm dish. Individual clones were harvested and expanded. PCR products from each clone were sequenced and protein expression was confirmed by immunoblotting assay.

#### ER-reporter plasmid transfection and stable cell line generation

HEK293T cells were dissociated with 0.25% Trypsin and seeded at a density of 30,000/cm2. Cells were transduced with lentiviruses expressing the ER-phagy reporter (Addgene #128257) during splitting in MEF medium. Stable transformants were selected with puromycin (1 μg/ml). HEK293T cells stably expressing the ER-phagy reporter were cultured with 2 μg/mL Doxycycline for 24 h. Thereafter, cells were washed with PBS twice and incubated in Glucose-free DMEM (D6546; Sigma-Aldrich) supplemented with 10% FBS (S1820-500; Biowest), 2 mM L-glutamine (25030-081; GIBCO) followed by confocal microscopy or western blotting assay.

#### ER-phagy reporter transfection in human iNeurons

Mature human iNeurons (Day 42) were transfected with ER-phagy reporter plasmids using the JetOptimus Transfection kit. 48 h post transfection, the medium was changed to Glucose and Sodium pyruvate-free NBA (Gibco, A24775-01) supplemented with B27 (Life Technologies, 17504044), GlutaMAX (Gibco, 35050061), 1% dialyzed Fetal Bovine Serum (Gibco, A3382001) for 48h and fixed with 4% PFA at room temperature for 10 minutes and washed with PBS buffer for 3 times. Cells were mounted with Antifade Mounting Medium with DAPI (Vector Laboratories, Cat# NC9524612, 0.5 μg5ug/ml) directly then imaged by fluorescence microscopy.

#### Fluorescence microscopy of ER-phagy reporter

WT and *ATG7* KD human iNeuronswere cultured on coverslips coated with GelTrex. RFP-GFP-KDEL plasmids were delivered to WT and *ATG7* KD human iNeurons via JetOptimus transfection reagent. Autophagy was induced in human iNeurons by glucose and sodium pyruvate deprivation in Glucose and Sodium pyruvate-free NBA medium (Gibco, A2477501) supplemented with dialyzed FBS for 48h. Upon the completion of the nutrient starvation, neurons were fixed with 4% PFA at room temperature for 10 minutes and washed with TBST. Cells were mounted with Antifade Mounting Medium with DAPI (Vector Laboratories, Cat# NC9524612, 0.5 μg5ug/ml) directly then imaged by fluorescence microscopy and processed in ImageJ and Adobe Photoshop.

#### Lentivirus production

Lentiviruses were produced as described(Marro and Yang, 2014) in HEK293T cells (ATCC, passage number lower than 20) by co-transfection with three helper plasmids (pRSV-REV, pMDLg/pRRE and vesicular stomatitis virus G protein expression vector) using Polyethylenimine (PEI) (Longo et al., 2013). Lentiviral particles were ultra-centrifuged at 21,000rpm, 4°C for 2h, re-suspended in DMEM, aliquoted, and stored at −80°C. Only virus preparations with > 90% infection efficiency as assessed by EGFP expression or puromycin resistance were used for experiments

#### Generation of neurons from human PSCs

Glutamatergic neurons were generated by overexpression of the transcription factor Ngn2 as previously described(Zhang et al., 2013). Briefly, human PSCs were dissociated using 0.5 mM EDTA/PBS and plated at a density of 88,000 cells/cm2 in N2 medium supplemented with 50 mM Chroman1. At the same time, cells were mixed with FUW-TetO-Ngn2-P2A-puromycin and FUW-rtTA lentiviruses by adding them directly to the medium. 24 h later, the medium was replaced with N2 medium (1×N2 supplement, 1×NEAA in DMEM-F12 medium) containing Doxycycline (2mg/mL) to induce transgene expression. Transduced cells were enriched with puromycin (1mg/mL) for 2 days. 4-5 days post Doxycycline, neurons were dissociated and plated together with mouse glial cells (100,000 cells/cm2) on Geltrex-coated plates. Two weeks after transgene induction, Doxycycline was removed and the neuronal culture was maintained in Neurobasal A medium supplemented with 1×B27, 1×Glutamax, and 1% fetal bovine serum. Mature neurons were used for various experiments on day 42 (after 5 weeks of co-culture with mouse glia).

#### RNA extraction and mRNA sequencing

To determine gene expression level, *ATG7* KD and Control iNeurons (Day42, 1 well of a 6-well plate around 2.5×105 cells total) were washed with PBS and lysed in 1 mL of Trizol for RNA extraction. Following lysis, DNA contamination was removed using TURBO DNA-free kit treatment. 3 biological replicates were obtained for both *ATG7* KD and Control iNeurons.

Three pairs of mouse whole brain tissue from *Atg7*f/f and *Atg7*f/f-SynCre mice were dissected and snapped frozen using liquid nitrogen. Mice whole brains were grinded using the Mortar & Pestle Set (Thermofisher, Cat# FB961N and 50-195-4054054).

#### Western blot analysis of cell lysate

Cultured cells were harvested and were subjected to western blot. Cells were lysed in lysis buffer (1% Triton-X 100, 50 mM Tris HCl (pH=7.5), 150 mM NaCl, proteinase/phosphatase inhibitor, and EDTA) and supernatants were collected on ice. Protein concentrations were measured by the Pierce BCA Protein Assay Kit in accordance with the manufacturer’s protocol. 25 μg of total proteins were run on 4%-12% 15 well Bis-Tris gel and proteins were transferred to a PVDF membrane. Resulting membranes were blocked in a blocking buffer (5% non-fat milk in TBST or LI-COR Blocking Buffer (LI-COR, 927-60001) buffer for 1 hour at room temperature. Primary antibodies were diluted in blocking buffer as described above and incubated at 4°C for 24 h. Secondary antibodies were diluted in blocking buffer and incubated for 1 hour at room temperature. Membranes were visualized and processed with Image Lab and Adobe Photoshop.

#### Western blot analysis of mouse brain lysate

Whole brains were collected from mice described above at 6-8 week of age and homogenized with a lysis buffer containing 50 mM Tris HCl, pH 7.5, 150 mM NaCl, 1% Triton X-100, and proteinase/phosphatase inhibitor. Brain lysates were subjected to immunoblotting assay following the same protocol for cultured cells.

#### Immunofluorescence staining in cultured cells

Coverslips were washed once with PBS and then fixed with 4% PFA at room temperature for 10 minutes and with methanol (prechilled in -20°C) in -20°C for 30 minutes. Cells were then permeabilized with 0.2% Triton-X-100 for 10 minutes. The cells were then blocked with blocking buffer (1% CCS, 4% BSA in PBS) for 1-2 hours at room temperature. Primary antibodies diluted in blocking buffer containing 0.03% Triton-X-100 were incubated overnight at 4°C and secondary antibodies diluted in PBS containing 0.03% Triton-X-100 were incubated at room temperature for 1 hour. The cells were imaged under confocal microscopy at 40x and 63x and the figures were processed with ImageJ and Adobe Photoshop.

#### Immunofluorescence staining in mice brains

Mouse brains (6-8 weeks old) were perfused with 4% PFA for 10 minutes. They were fixed overnight in 4% PFA and 30% sucrose for 2 days at 4°C. After removal of sucrose, brains were placed in OCT compounds for cryosection and then incubated overnight at -80°C. The block was transferred to the cryostat 30 minutes before cutting and incubated, and then sections were cut into 30 m thick at -20°C. These slices were transferred to 24-well plate in PBS. Tissues were then permeabilized with 0.3% Triton-X-100 for 20 minutes and blocked with blocking solution consisting of 5% goat serum and 0.3% Triton-X-100 for 1 hour at room temperature. Primary antibodies were incubated overnight at 4°C and secondary antibodies were incubated at room temperature for 1 hour. Imaging was then performed with a confocal microscopy on 10x, 20x, and 63x objective using Z-stack and tile scan tools and analyzed with image J.

#### Immunoprecipitation

HEK293T cells were lysed in IP buffer (1% NP-40, 10 mM Tris-HCl pH 7.5, 100 mM NaCl, 2 mM EDTA, 1xHalt™ protease and phosphatase inhibitor cocktail (Thermofisher, Cat# WE325176)) for 30 minutes on ice and centrifuged at 13,000 g for 30 minutes at 4°C. Supernatants were collected after centrifugation and subjected to BCA assay. Supernatants containing 375 µg of total protein were incubated with GFP-Trap Magnetic Agarose (Chromotek, Cat# 010-272-01-01) antibodies overnight at 4°C. GFP-Trap Magnetic Beads was washed with PBS (0.1% Triton X-100) 3 times, 5 minutes each, 1X SDS loading buffer was added to the beads and boiled at 70 □ for 10 minutes. The eluate was collected and subjected to Western Blot. Total protein of 10 µg10ug was loaded for input.

#### Screening strategy for ideal candidate gene from autophagy-deficient human iN MS data

A multi-step screening process was used to determine the most ideal candidate gene for our study. The first step in establishing a list of candidate genes was to screen for genes in the *ATG7* and *ATG14* KD human iNeurons proteomics data that had a Log2FC greater than the standard deviation (Log2FC>SD) and a Log2FC greater than 0 (Log2FC>0). These genes were then subjected to GO:TERM analysis by a web-based tool to determine which genes were related to the ER (https://biit.cs.ut.ee/gprofiler/gost). SynGo was performed in the webtool (https://www.syngoportal.org/). Genes that were related to the ER were next screened by a web-based tool designed to screen a protein’s amino acid sequence for the presence of a conserved LIR motif (https://ilir.warwick.ac.uk/search.php). From this web-based tool, proteins that had LIR motifs in their ANCHOR regions (intrinsically disordered region) were selected. The remaining proteins were screened with a third web-based tool that predicts the presence of a transmembrane domain in a protein’s secondary structure (http://bioinf.cs.ucl.ac.uk/psipred/). Proteins that satisfied all three prerequisites were included in the final list of candidate proteins.

#### Electron microscopy (EM)

Three pairs of *Atg7*f/f and *Atg7*f/f-SynCre (*Atg7* cKO) mice were sacrificed, and samples were processed following the procedures described in(Wang et al., 2006). Similarly, 3 pairs of *Beclin1*f/f and *Beclin1*f/f-P2pCre mouse brain samples were processed as described in(McKnight et al., 2014).

#### Sample preparation and quantitative proteomics Autophagy-deficient human iNeurons and mouse brains

Three batches of H1-Lenti-*ATG*7/14 KD derived Ngn2 iNeurons were included for the quantitative proteomics and phosphoproteomics analyses (day 42, two subclones for each genotype. ATG7 KD: # 1, #6; ATG14 KD: #3, #5; Vector: #2, #6). The cells were washed with PBS once and manually dissociated by gentle trituration in PBS. Cells were then centrifuged at 500g for 3 minutes. Supernatant was removed, and cell pellet was snap froze by liquid nitrogen and stored in -80°C before subject to proteomics analyses. For mouse brain samples, 6-8-week-old autophagy-deficient mice (3 *Atg7*f/f-SynCre and 4 *Atg7*f/f mice; 3 *Atg14*f/f-SynCre and 4 *Atg14*f/f mice) were sacrificed and whole brains were dissected, and snap frozen in liquid nitrogen and stored in -80°C before proteomics analyses.

#### Protein extraction, quantification of iNeurons and mouse brain samples

iN cell pellets and mouse brain samples were lysed, and the protein concentrations were quantified as previously described (Bai et al., 2017). Briefly, the frozen samples were homogenized in the lysis buffer (50 mM HEPES, pH 8.5, 8 M urea, and 0.5% sodium deoxycholate) with 1X PhosSTOP phosphatase inhibitor cocktail (Sigma-Aldrich). Protein concentration was measured by the BCA assay (Thermo Fisher) and then confirmed by Coomassie-stained short SDS gels (Xu et al., 2009).

#### Protein digestion and TMT labeling for iNenrons and mouse brain samples

The analysis was performed with a previously optimized protocol (Bai et al., 2020) ∼0.1 mg of iNeurons protein samples and ∼0.1 mg of mouse brain protein samples (in lysis buffer with 8 M urea) were first digested by Lys-C (Wako, 1:100 w/w) at 21□°C for 2 h and then diluted by 4-fold to reduce urea to 2 M by 50mM HEPES followed by the addition of trypsin (Promega, 1:50 w/w) to continue the digestion at 21°C overnight. The digestion was terminated by the addition of 1% trifluoroacetic acid. After centrifugation, the supernatant was desalted and then dried by Speedvac. Each sample was resuspended in 50 mM HEPES (pH 8.5) for TMT labeling, and then mixed equally followed by desalting. The iN samples were labeled by TMT16. The *Atg7* cKO and *Atg14* cKO mouse brain samples were labeled by two sets of TMT10 respectively, but only 7 TMT channels was discussed here.

#### Phosphopeptide enrichment of iN samples

Phosphopeptide enrichment was carried out by TiO_2_ beads as descripted previously (Tan et al., 2015). Briefly, the pooled desalted TMT labeled peptides (∼1.8 mg) were dissolved in the binding buffer (65% acetonitrile, 2% TFA, and 1 mM KH2PO4). TiO_2_ beads (7.2 mg) were washed twice with the washing buffer (65% acetonitrile, 0.1% TFA), incubated with the peptide at 21 °C for 20 minutes. The TiO_2_ beads were then washed twice again with washing buffer and loaded into a C18 StageTip (Thermo Fisher), followed by the elution of phosphopeptides by the basic pH buffer (15% NH4OH, and 40% acetonitrile). The eluates were dried respectively and dissolved in 5% formic acid for LC-MS/MS analysis. The flowthrough of phosphopeptide enrichment was desalted and dried for basic LC fractionation.

#### GFP-LC3 affinity purification from mouse brains, in-gel digestion, and TMT labeling

The GFP-LC3 overexpression and control mouse brains were harvested and rinsed with cold PBS. Homogenization buffer A (0.32 M sucrose, 1 mM NaHCO3, 0.25 mM CaCl2, 1 mM MgCl2, 50 mM Tris HCl, pH 7.5) supplied with halt proteinase and phosphatase inhibitors (78442, Invitrogen) was added as 2 ml buffer A per mouse brain. The tissue was homogenized with a glass douncer 20 times before being centrifuged at 1000g for 10 minutes. The supernatant was transferred to a new 15-ml falcon tube labeled as the cytoplasmic sample. 2X buffer D (300 mM NaCl, 20% NP40, 10% Sodium Deoxycholate, 2 mM EDTA, 2 mM EGTA, 100 mM Tris HCl, pH 7.5) was added and the sample was rotated for 1 h. Next, the sample was centrifuged at 20,800g for 20 minutes and the supernatant was transferred into a new falcon tube. The sample was then centrifuged for the second time using the same conditions, with the supernatant being collected. 100 µl was saved as input and protein concentration was quantified. 15µl GFP-trap beads (gmta-20 GFP-Trap ChromoTek GmbH) per mouse brain was added and the sample was incubated overnight. The next day, beads were washed three times (10 minutes each, 500 µl of 1X buffer D). After the final wash, the protein was eluted at 70°C with 30 µl of 1X LDS-PAGE sample buffer (NP0007). 3 µl of the IP sample was used for SDS-PAGE and Silver staining (LC6070, Invitrogen). The rest of the IP samples were run on a very short SDS gel followed by in-gel digestion(Bai et al., 2013), the peptides were then labeled by TMT11 respectively, mixed equally, followed by desalting for the subsequent fractionation. Only 8 of the 11 channels’ data was discussed here.

#### Extensive two-dimensional liquid chromatography-tandem mass spectrometry

For each of the 4 TMT sets (iNeurons, *Atg7* cKO mouse brain, *Atg14* cKO mouse brain and GFP-LC3 affinity purification samples), the TMT labeled samples were fractionated by offline basic pH reverse phase LC respectively, and each of these fractions was analyzed by the acidic pH reverse phase LC-MS/MS(Niu et al., 2017). The offline basic pH LC was performed with an XBridge C18 column (3.5□μm particle size, 4.6□mm□×□□25□cm, Waters), buffer A (10□mM ammonium formate, pH□8.0), buffer B (95% acetonitrile, 10□mM ammonium formate, pH□8.0), using a 2–3□h gradient of 15–35% buffer B (Bai et al., 2017). In the acidic pH LC-MS/MS analysis, fractions were analyzed sequentially on a column (75□μm×15–30□cm, 1.9□μm C18 resin from Dr. Maisch GmbH, 65□°C to reduce backpressure) coupled with a Fusion or Q Exactive HF Orbitrap mass spectrometer (Thermo Fisher Scientific). Peptides were analyzed with a 1–3□h gradient (buffer A: 0.2% formic acid, 5% DMSO; buffer B: buffer A plus 65% acetonitrile). For mass spectrometer settings, positive ion mode and data-dependent acquisition were applied with one full MS scan followed by a 20 MS/MS scans. MS1 scans were collected at a resolution of 60,000,1□×□10^6^ AGC and 50□ms maximal ion time; higher energy collision-induced dissociation (HCD) was set to 32–38% normalized collision energy; ∼□1.0□m/z isolation window with 0.3□m/z offset was applied; MS2 spectra were acquired at a resolution of 60,000, fixed first mass of 120□□/z, 410–1600□m/z, 1□×□10^5^ AGC, 100–150□ms maximal ion time, and□~□15□s of dynamic exclusion. For the phosphoproteomics analysis, 1.5 m/z isolation window was used to improve sensitivity.

#### Protein and phosphopeptide identification and quantification by the JUMP software suite

The bioinformatics processing of protein and phosphopeptide identification and quantification were carried out with the JUMP software suites(Wang et al., 2014). In brief, MS/MS raw data were searched against a target-decoy database to estimate false discovery rate (FDR)(Peng et al., 2003). We combined the downloaded Swiss-Prot, TrEMBL, and UCSC databases and removed redundancy (human: 83,955 entries; mouse, 59,423 entries) to create the database. Main search parameters were set at precursor and product ion mass tolerance (±15□ppm), full trypticity, maximal modification sites (n□=□3), maximal missed cleavage (n□=□2), static mass shift including carbamidomethyl modification (+□57.02146 on Cys), TMT tags (+u□229.16293 for TMT10/11 or + 304.20714 for TMT16 on Lys and N-termini), and dynamic mass shift for oxidation (+□15.99491 on Met) and Ser/Thr/Tyr phosphorylation (+79.96633). Peptide-spectrum matches (PSM) were filtered by mass accuracy, clustered by precursor ion charge, and the cutoffs of JUMP-based matching scores (J-score and ΔJn) to reduce FDR below 1% for proteins during the whole proteome analysis or 1% for phosphopeptides during the phosphoproteome analysis. Protein and phosphopeptide quantifications were performed based on the reporter ions from MS2 using our previously optimized method(Niu et al., 2017).

#### Quantification and Statistical Analysis

Unless otherwise indicated, all data presented are the average of at least two biological replicates from each of at least two independent experiments. See figure legends for details on specific statistical tests run for each experiment. Error bars were calculated using the standard deviation of all the replicates. Statistical significance is represented by a star (∗ = p < 0.05, ∗∗ = p < 0.005 and ∗∗∗ = p < 0.0005, ∗∗∗∗= p < 0.0001).). Graphs and plots were generated using Graphpad Prism 9.0 or RStudio (2022.07.1). Figures were generated using Adobe Illustrator 2022.

## Supplemental information

**Figure S1.**
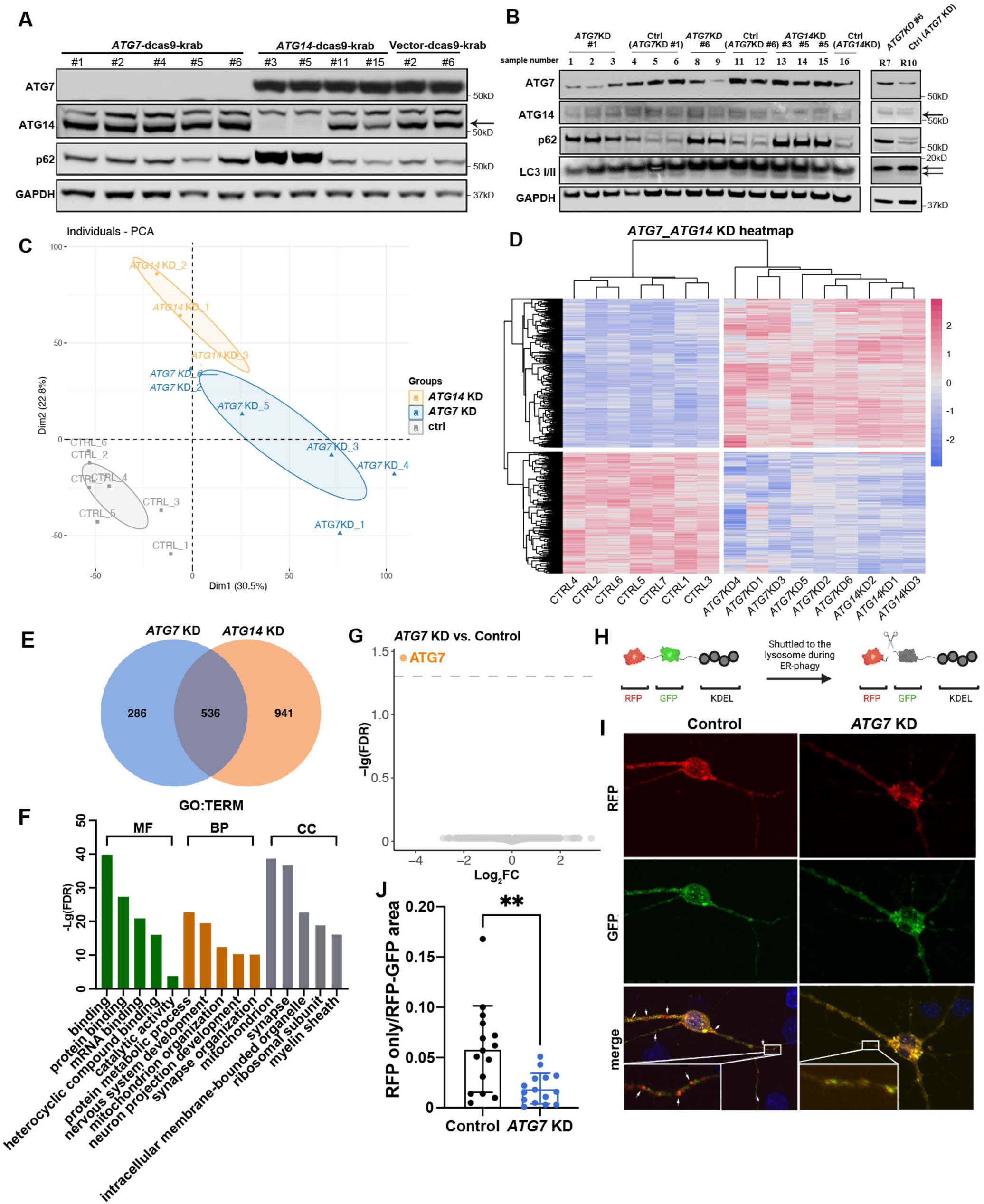
Generation and proteomic analysis of human *ATG7* or *ATG14* KD iNeurons. (A) Immunoblot analysis of multiple clones of *ATG7* KD, *ATG14* KD, or control with antibodies as indicated. Human mutant iNeurons generated from subclones *ATG7* KD#1 and #6, *ATG14* KD#3 and #5, Vector #2 and #6 were selected for proteomics analysis. (B) Immunoblot analysis of indicated iNeuron clone samples submitted for proteomic analysis. (C) Principal component analysis (PCA) of 6 *ATG7* KD iNeurons (blue), 3 *ATG14* KD iNeurons (orange), and 6 Control iNeurons (gray) for proteomics. (D) Heatmap analysis of Log2FC of DEPs (FDR<0.05) for human iN proteomics samples. (E) Venn diagram showing the overlap between the downregulated DEPs of *ATG7* KD (blue) and of *ATG14* KD (orange) iNeurons (FDR< 0.05). (F) GO enrichment analysis of 536 downregulated DEPs shared between *ATG7* KD and *ATG14* KD iNeurons from (E) (FDR< 0.05). MF: Molecular Function, BP: Biological Process, CC: Cellular Component. (G) Volcano plot of the DEGs from transcriptomic data of three batches of *ATG7* KD iNeurons vs. Control iNeurons. Orange dots represent the only DEG identified (FDR< 0.05). Dashed line is at -Lg (FDR)= 1.3. (H) Schematic of the ER-phagy reporter RFP-GFP-KDEL. The GFP signal is quenched when fused with lysosomes to yield the RFP only signal. (I-J) Immunofluorescence images (I) of Control and *ATG7* KD iNeurons transfected with ER-phagy reporter plasmids after nutrient starvation (glucose and sodium pyruvate starvation for 48 h and quantification of the percentage of RFP only area (J). Inset, magnified images of the boxed μm. N=15, unpaired t-test. **p<0.01.

**Figure S2.**
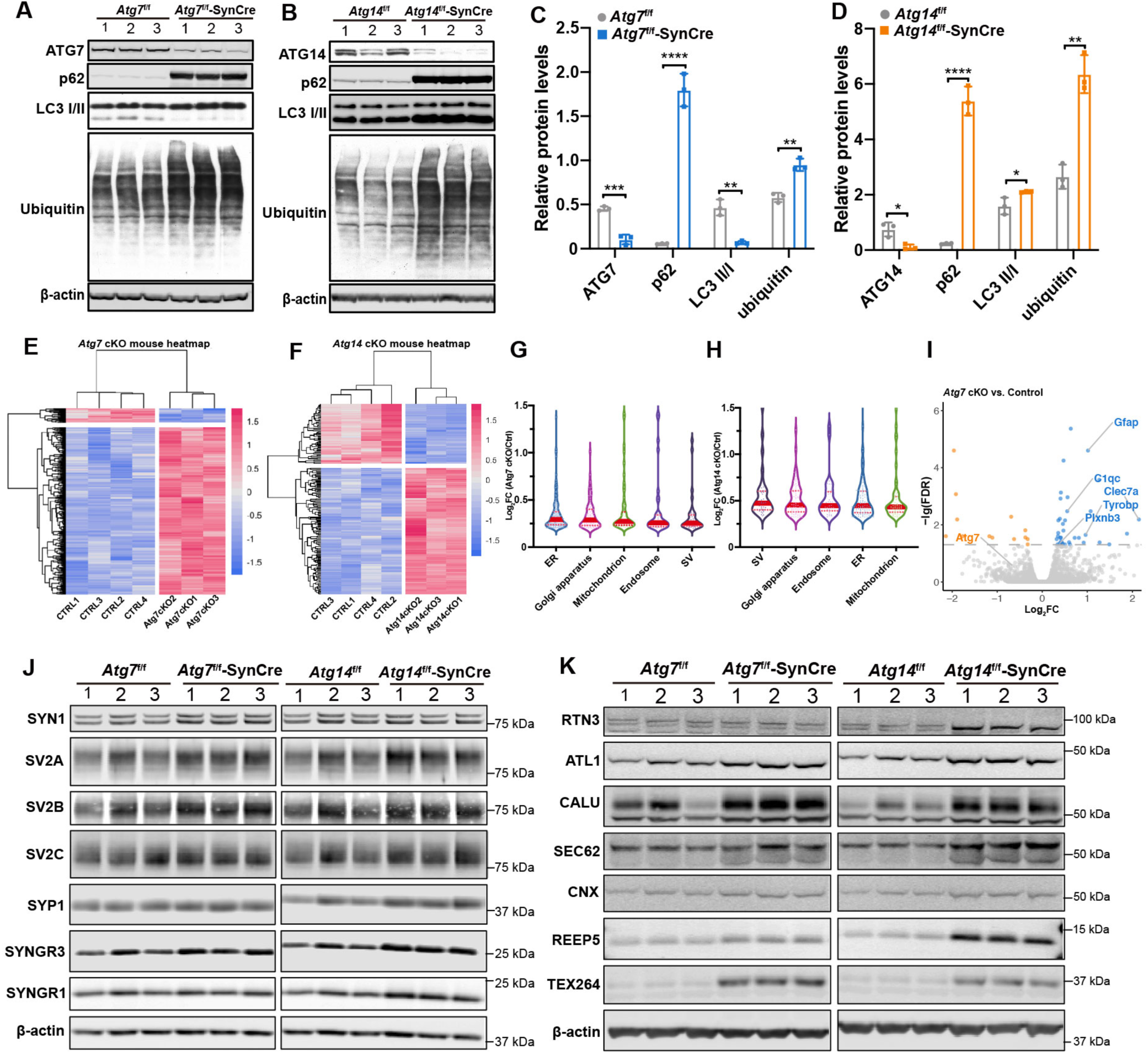
Proteomic and transcriptomic analyses of *Atg7* or *Atg14* cKO mouse brains. (A-B) Immunoblot analysis of indicated autophagy marker proteins in *Atg7*f/f and *Atg7*f/f-SynCre (A); *Atg14*f/f and *Atg14*f/f-SynCre (B) mouse brains (2-months old), n= 3. (C-D) Quantification of the change of indicated autophagy marker proteins from (A) and (B). Relative protein levels were normalized to the loading control β actin. Unpaired t-test. *p<0.05, **p<0.01, ***p<0.001, ****p<0.001. (E-F) Heatmap analysis of Log2FC of DEPs (*p* < 0.05, | Log2FC |>2SD) for *Atg7* cKO (E) and *Atg14* cKO (F) mouse proteomics samples. (G and H) Violin plots of DEPs in ER, SV, Golgi apparatus, mitochondria, and endosome based on the GO analysis in (Figure 3E) and (Figure 3F), respectively. Each dot represents one protein. Solid red bars indicate the median Log2FC, and the red dashed bars specify the 25th and 75th interquartile range. (I) Volcano plot of the DEGs from transcriptomic data of *Atg7*f/f and *Atg7*f/f-SynCre mouse brains (n=3). Blue dots represent upregulated genes (FDR< 0.05, Log2FC> 0) and orange dots represent downregulated genes. Dashed line is at -Lg (FDR)= 1.3. (J) Immunoblot analysis of ER proteins as indicated in *Atg7*f/f and *Atg7*f/f-SynCre (left); *Atg14*f/f and *Atg14*f/f-SynCre (right) mouse brains (2-month-old) (n= 3). (K) Immunoblot analysis of SV proteins as indicated in *Atg7*f/f and *Atg7*f/f-SynCre (left); *Atg14*f/f and *Atg14*f/f-SynCre (right) mouse brains (2-month-old) (n= 3).

**Figure S3.**
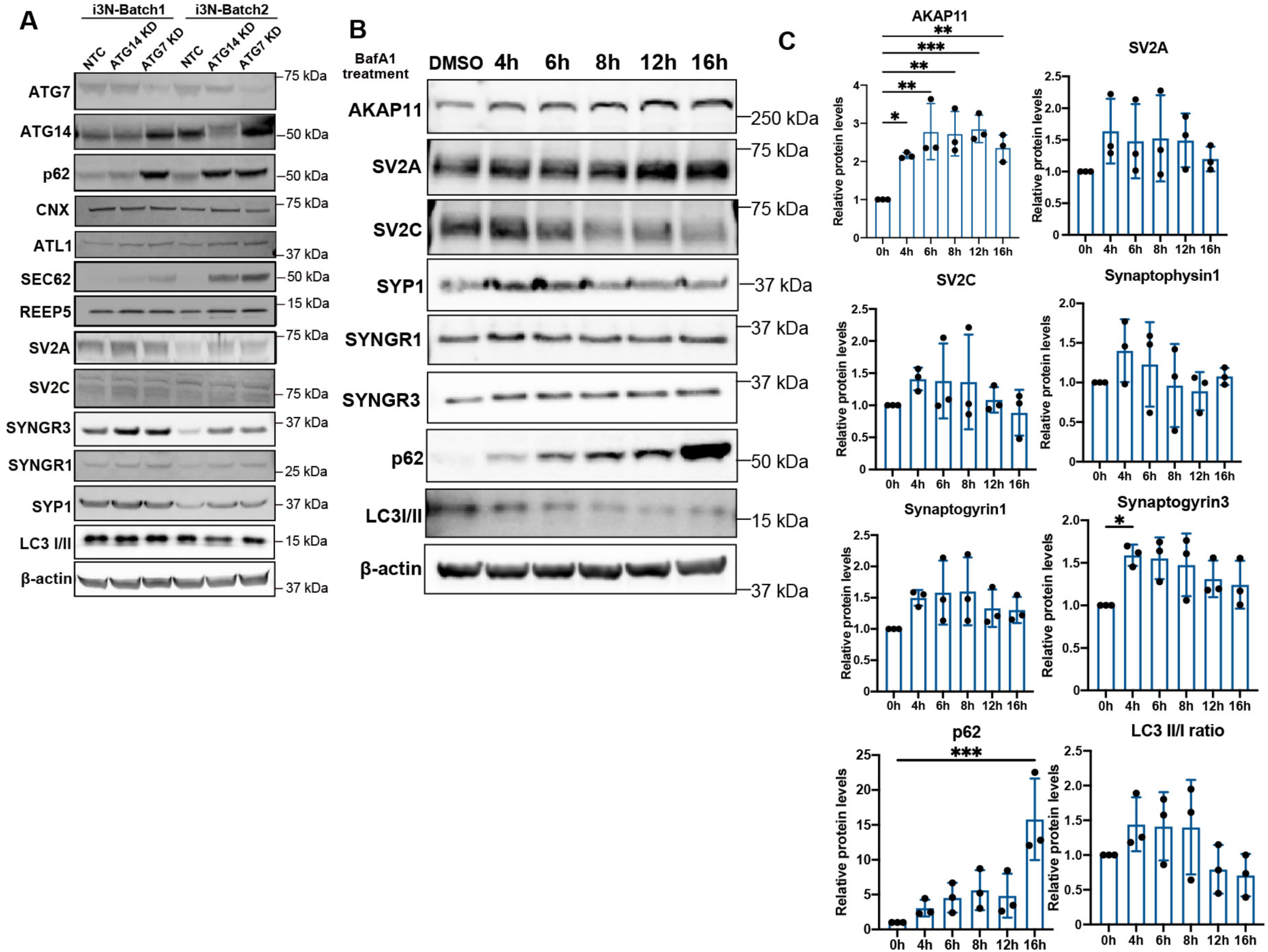
Analysis of ER and SV related proteins in multiple iNeuron lines. (A) Immunoblot analysis of ER and SV proteins as indicated in control, *ATG7* KD, and *ATG14* KD iNeurons derived from the inducible iPSC cell lines (i3N) from two independent biological replicates. (B and C) Immunoblot analysis of ER and SV proteins as indicated in WT iNeurons upon a time-dependent (4h, 6h, 8h, 12h, 16h) treatment of Bafilomycin A1(100nM). DMSO-treated iNeurons serve as a negative control. Relative protein levels were normalized to the loading control β actin (C). Unpaired t-test, three replicates. *p<0.05, **p<0.01, ***p<0.001, ****p<0.001.

**Figure S4.**
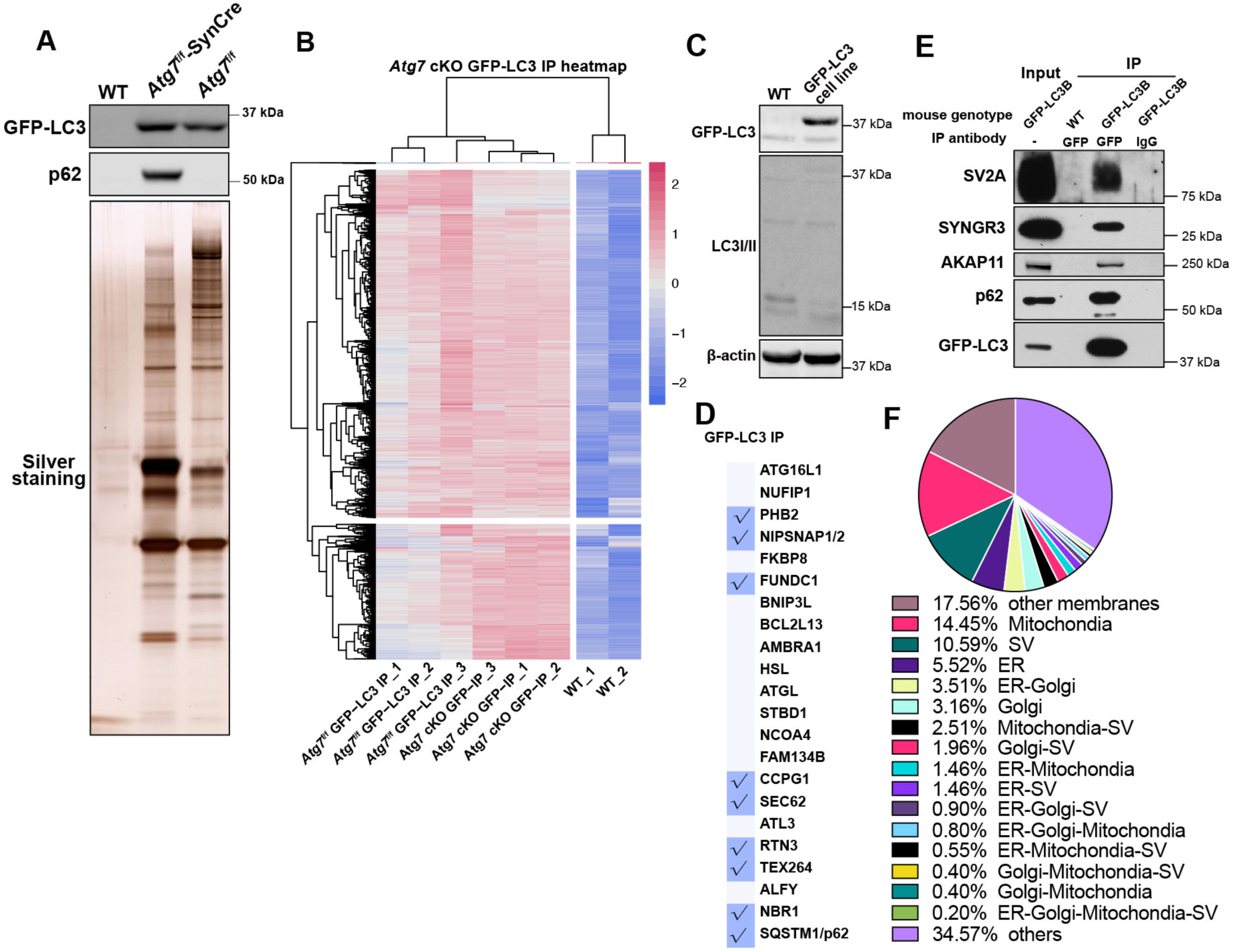
Isolation and detection of LC3-interacting proteins from mouse brains. (A) Protein chemistry study of indicated proteins for *GFP-LC3*; *Atg7*f/f or *GFP-LC3*; *Atg7*f/f-SynCre mouse brain samples subjected to GFP-LC3 affinity purification and subsequent proteomics. Top and middle panels, immunoblot analysis with indicated antibodies. Bottom, silver staining of proteins after GFP-LC3 affinity purification. (B) Heatmap analysis of Log2FC of DEPs (*p* < 0.05, | Log2FC |>2SD) for samples from GFP-LC3 affinity purification proteomics. (C) Immunoblot analysis confirms the endogenous LC3 and GFP-LC3 proteins in the human WTC11 iPSC cells line stably expressing GFP-LC3. (D) A table of the known autophagy receptors and those detected (check marks) in proteomic analysis of *Atg7*f/f-SynCre; GFP-LC3 IP (*p* < 0.05, | Log2FC |>2SD, Log2FC> 0). The list of autophagy receptors was manually collated through literature searches. (E) Co-immunoprecipitation and immunoblot analysis of the indicated proteins through GFP-LC3 affinity pull-down from GFP-LC3 transgenic and WT mouse brains. (F) Pie chart analysis of the percentage of proteins categorized into different organelles or pathways from upregulated DEPs identified *GFP-LC3*; *Atg7* cKO IP compared to the WT control (*p* < 0.05, | Log2FC |>2SD, Log2FC> 0). Categories were summarized from GO term analysis.

**Figure S5.**
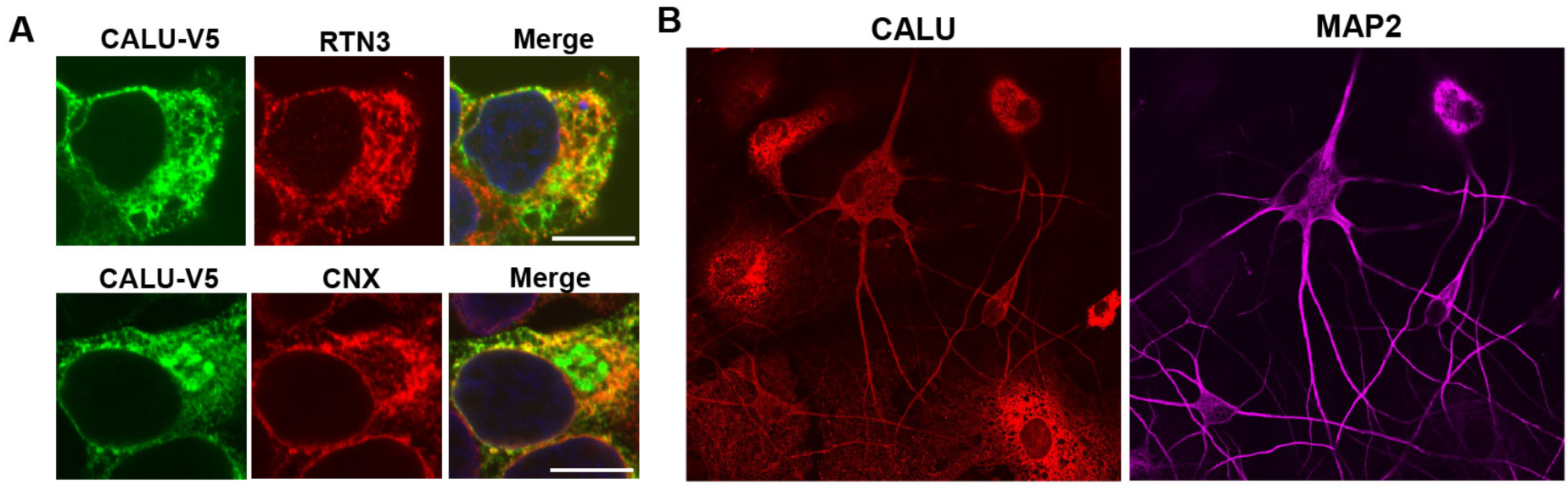
Characterization of CALU localization and CALU KO cells. (A) Immunofluorescence images of HEK293T cells transfected with CALU-v5 plasmids and co-stained with anti-v5 (green) and anti-RTN3 or anti-CNX (Calnexin) antibodies. Scale bar, 10 μm. (B) Immunofluorescence images of human iNeurons (6-week-old) co-stained with anti-CALU (red) and anti-MAP2 (green) antibodies. Scale bar, 10 μm.

**Figure S6.**
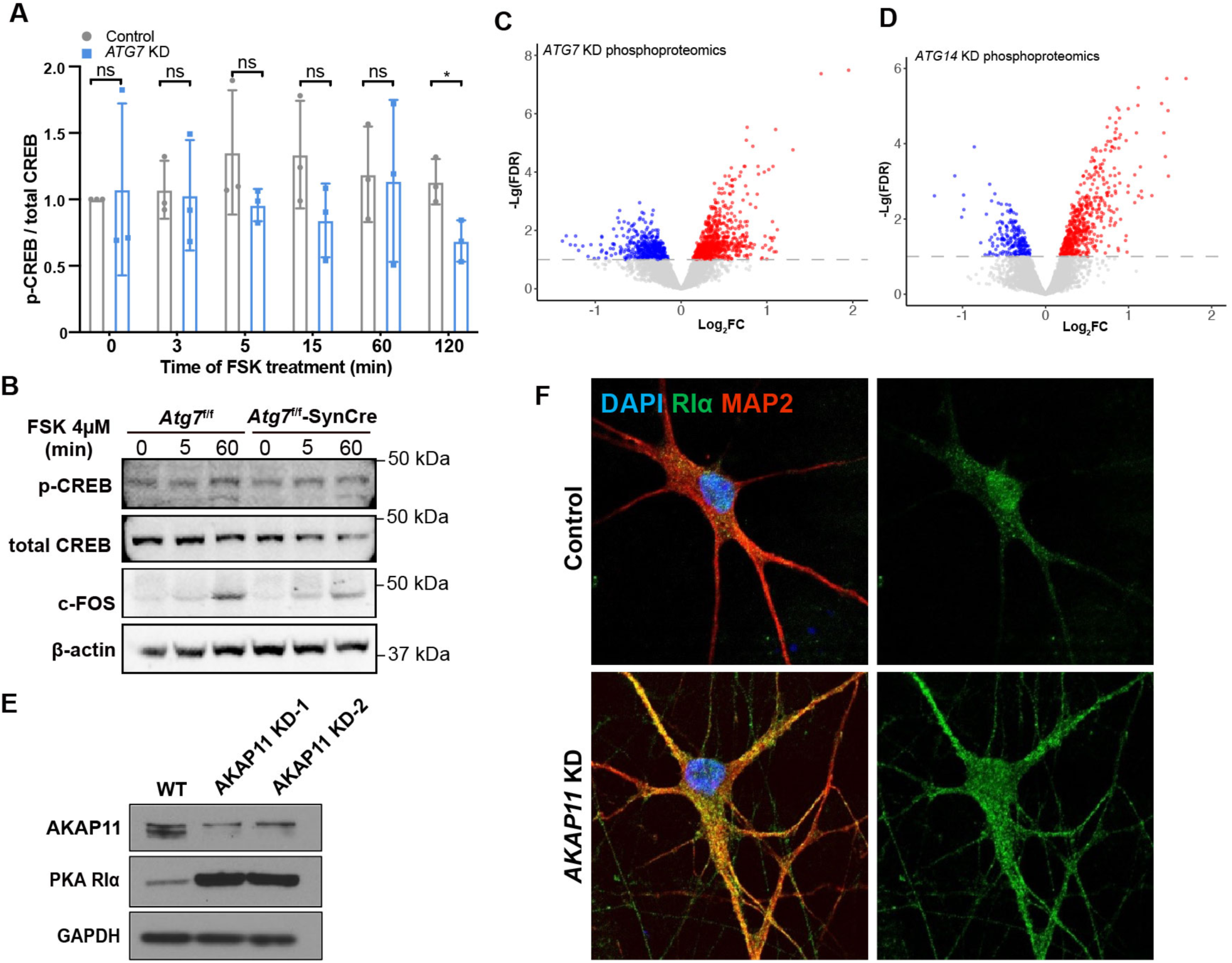
Analysis of PKA kinase activity in *ATG7* KD iNeurons and mouse primary neurons from *Atg7* cKO mice. (A) Quantification for (Figure 6D) of p-CREB normalized to total CREB from in control and *ATG7* KD human iNeurons from i3N-derived iNeurons treated with Forskolin (4 μM) for 3, 5, 15, 60, 120minutes. Relative protein levels were normalized to β-actin. Data were collected from three biological replicates. Un-paired *t*-test. *p<0.05, ns, no significance. (B) Immunoblot analyses of indicated proteins in primary cortical neurons (DIV14) cultured from *Atg7*f/f and *Atg7*f/f-SynCre mice after time-dependent (0 h, 5minutes, 1 h) Forskolin treatment (4 μM). Representative images from two independent replicates. (C-D) Volcano plots of differentially phosphorylated proteins identified from phosphoproteomic analyses in *ATG7* KD (left) and *ATG14* KD (right) iNeurons. Red and blue dots represent proteins with an increased phosphorylation (FDR< 0.05, Log2FC> 0) and decreased phosphorylation (FDR< 0.05), respectively. Dashed line is at -Lg (FDR)= 1.3. (E) Immunoblot analysis of AKAP11 and PKA-RΙα protein levels in two clones of *AKAP11* KD human iNeurons. (F) Immunofluorescence images of Control and *AKAP11* KD iNeurons.

## Notes

### Competing Interest Statement

The authors have declared no competing interest.

## References

An, H., Ordureau, A., Paulo, J.A., Shoemaker, C.J., Denic, V., and Harper, J.W. (2019). TEX264 Is an Endoplasmic Reticulum-Resident ATG8-Interacting Protein Critical for ER Remodeling during Nutrient Stress. Mol Cell 74, 891–908 e810.

Bai, B., Hales, C.M., Chen, P.C., Gozal, Y., Dammer, E.B., Fritz, J.J., Wang, X., Xia, Q., Duong, D.M., Street, C., et al. (2013). U1 small nuclear ribonucleoprotein complex and RNA splicing alterations in Alzheimer’s disease. Proc Natl Acad Sci U S A 110, 16562–16567.

Bai, B., Tan, H., Pagala, V.R., High, A.A., Ichhaporia, V.P., Hendershot, L., and Peng, J. (2017). Deep Profiling of Proteome and Phosphoproteome by Isobaric Labeling, Extensive Liquid Chromatography, and Mass Spectrometry. Methods Enzymol 585, 377–395.

Bai, B., Wang, X., Li, Y., Chen, P.C., Yu, K., Dey, K.K., Yarbro, J.M., Han, X., Lutz, B.M., Rao, S., et al. (2020). Deep Multilayer Brain Proteomics Identifies Molecular Networks in Alzheimer’s Disease Progression. Neuron 105, 975–991 e977.

Bampton, E.T., Goemans, C.G., Niranjan, D., Mizushima, N., and Tolkovsky, A.M. (2005). The dynamics of autophagy visualized in live cells: from autophagosome formation to fusion with endo/lysosomes. Autophagy 1, 23–36.

Behrends, C., Sowa, M.E., Gygi, S.P., and Harper, J.W. (2010). Network organization of the human autophagy system. Nature 466, 68–76.

Birgisdottir, A.B., Lamark, T., and Johansen, T. (2013). The LIR motif - crucial for selective autophagy. J Cell Sci 126, 3237–3247.

Chen, Q., Xiao, Y., Chai, P., Zheng, P., Teng, J., and Chen, J. (2019). ATL3 Is a Tubular ER-Phagy Receptor for GABARAP-Mediated Selective Autophagy. Curr Biol 29, 846–855 e846.

Chino, H., Hatta, T., Natsume, T., and Mizushima, N. (2019). Intrinsically Disordered Protein TEX264 Mediates ER-phagy. Mol Cell 74, 909–921.e906.

Collier, J.J., Guissart, C., Oláhová, M., Sasorith, S., Piron-Prunier, F., Suomi, F., Zhang, D., Martinez-Lopez, N., Leboucq, N., Bahr, A., et al. (2021). Developmental Consequences of Defective ATG7-Mediated Autophagy in Humans. N Engl J Med 384, 2406–2417.

Day, M.E., Gaietta, G.M., Sastri, M., Koller, A., Mackey, M.R., Scott, J.D., Perkins, G.A., Ellisman, M.H., and Taylor, S.S. (2011). Isoform-specific targeting of PKA to multivesicular bodies. J Cell Biol 193, 347–363.

Deng, Z., Li, X., Blanca Ramirez, M., Purtell, K., Choi, I., Lu, J.H., Yu, Q., and Yue, Z. (2021). Selective autophagy of AKAP11 activates cAMP/PKA to fuel mitochondrial metabolism and tumor cell growth. Proc Natl Acad Sci U S A 118.

Dengjel, J., Høyer-Hansen, M., Nielsen, M.O., Eisenberg, T., Harder, L.M., Schandorff, S., Farkas, T., Kirkegaard, T., Becker, A.C., Schroeder, S., et al. (2012). Identification of autophagosome-associated proteins and regulators by quantitative proteomic analysis and genetic screens. Mol Cell Proteomics 11, M111.014035.

Dinkel, H., Chica, C., Via, A., Gould, C.M., Jensen, L.J., Gibson, T.J., and Diella, F. (2011). Phospho.ELM: a database of phosphorylation sites--update 2011. Nucleic Acids Res 39, D261–267.

Dubnau, J., Chiang, A.S., and Tully, T. (2003). Neural substrates of memory: from synapse to system. J Neurobiol 54, 238–253.

Friedman, L.G., Lachenmayer, M.L., Wang, J., He, L., Poulose, S.M., Komatsu, M., Holstein, G.R., and Yue, Z. (2012). Disrupted autophagy leads to dopaminergic axon and dendrite degeneration and promotes presynaptic accumulation of α-synuclein and LRRK2 in the brain. J Neurosci 32, 7585–7593.

Fumagalli, F., Noack, J., Bergmann, T.J., Cebollero, E., Pisoni, G.B., Fasana, E., Fregno, I., Galli, C., Loi, M., Solda, T., et al. (2016). Translocon component Sec62 acts in endoplasmic reticulum turnover during stress recovery. Nat Cell Biol 18, 1173–1184.

Geng, J., and Klionsky, D.J. (2017). Direct quantification of autophagic flux by a single molecule-based probe. Autophagy 13, 639–641.

Gesellchen, F., Bertinetti, O., and Herberg, F.W. (2006). Analysis of posttranslational modifications exemplified using protein kinase A. Biochim Biophys Acta 1764, 1788–1800.

Gold, M.G., Lygren, B., Dokurno, P., Hoshi, N., McConnachie, G., Tasken, K., Carlson, C.R., Scott, J.D., and Barford, D. (2006). Molecular basis of AKAP specificity for PKA regulatory subunits. Mol Cell 24, 383–395.

Goldsmith, J., Ordureau, A., Harper, J.W., and Holzbaur, E.L.F. (2022). Brain-derived autophagosome profiling reveals the engulfment of nucleoid-enriched mitochondrial fragments by basal autophagy in neurons. Neuron 110, 967–976.e968.

Grumati, P., Morozzi, G., Holper, S., Mari, M., Harwardt, M.I., Yan, R., Muller, S., Reggiori, F., Heilemann, M., and Dikic, I. (2017). Full length RTN3 regulates turnover of tubular endoplasmic reticulum via selective autophagy. Elife 6.

Hara, T., Nakamura, K., Matsui, M., Yamamoto, A., Nakahara, Y., Suzuki-Migishima, R., Yokoyama, M., Mishima, K., Saito, I., Okano, H., et al. (2006). Suppression of basal autophagy in neural cells causes neurodegenerative disease in mice. Nature 441, 885–889.

Hernandez, D., Torres, C.A., Setlik, W., Cebrian, C., Mosharov, E.V., Tang, G., Cheng, H.C., Kholodilov, N., Yarygina, O., Burke, R.E., et al. (2012). Regulation of presynaptic neurotransmission by macroautophagy. Neuron 74, 277–284.

Higuchi, H., Yamashita, T., Yoshikawa, H., and Tohyama, M. (2003). PKA phosphorylates the p75 receptor and regulates its localization to lipid rafts. EMBO J 22, 1790–1800.

Hockemeyer, D., Soldner, F., Cook, E.G., Gao, Q., Mitalipova, M., and Jaenisch, R. (2008). A drug-inducible system for direct reprogramming of human somatic cells to pluripotency. Cell Stem Cell 3, 346–353.

Jacomin, A.C., Samavedam, S., Promponas, V., and Nezis, I.P. (2016). iLIR database: A web resource for LIR motif-containing proteins in eukaryotes. Autophagy 12, 1945–1953.

Johansen, T., and Lamark, T. (2020). Selective Autophagy: ATG8 Family Proteins, LIR Motifs and Cargo Receptors. J Mol Biol 432, 80–103.

Kabeya, Y., Mizushima, N., Ueno, T., Yamamoto, A., Kirisako, T., Noda, T., Kominami, E., Ohsumi, Y., and Yoshimori, T. (2000). LC3, a mammalian homologue of yeast Apg8p, is localized in autophagosome membranes after processing. EMBO J 19, 5720–5728.

Khaminets, A., Behl, C., and Dikic, I. (2016). Ubiquitin-Dependent And Independent Signals In Selective Autophagy. Trends Cell Biol 26, 6–16.

Khaminets, A., Heinrich, T., Mari, M., Grumati, P., Huebner, A.K., Akutsu, M., Liebmann, L., Stolz, A., Nietzsche, S., Koch, N., et al. (2015). Regulation of endoplasmic reticulum turnover by selective autophagy. Nature 522, 354–358.

Kim, C., Cheng, C.Y., Saldanha, S.A., and Taylor, S.S. (2007). PKA-I holoenzyme structure reveals a mechanism for cAMP-dependent activation. Cell 130, 1032–1043.

Kirkin, V., Lamark, T., Johansen, T., and Dikic, I. (2009). NBR1 cooperates with p62 in selective autophagy of ubiquitinated targets. Autophagy 5, 732–733.

Komatsu, M., Waguri, S., Chiba, T., Murata, S., Iwata, J., Tanida, I., Ueno, T., Koike, M., Uchiyama, Y., Kominami, E., et al. (2006). Loss of autophagy in the central nervous system causes neurodegeneration in mice. Nature 441, 880–884.

Komatsu, M., Waguri, S., Koike, M., Sou, Y.S., Ueno, T., Hara, T., Mizushima, N., Iwata, J., Ezaki, J., Murata, S., et al. (2007a). Homeostatic levels of p62 control cytoplasmic inclusion body formation in autophagy-deficient mice. Cell 131, 1149–1163.

Komatsu, M., Waguri, S., Ueno, T., Iwata, J., Murata, S., Tanida, I., Ezaki, J., Mizushima, N., Ohsumi, Y., Uchiyama, Y., et al. (2005). Impairment of starvation-induced and constitutive autophagy in Atg7-deficient mice. J Cell Biol 169, 425–434.

Komatsu, M., Wang, Q.J., Holstein, G.R., Friedrich, V.L., Jr., Iwata, J., Kominami, E., Chait, B.T., Tanaka, K., and Yue, Z. (2007b). Essential role for autophagy protein Atg7 in the maintenance of axonal homeostasis and the prevention of axonal degeneration. Proc Natl Acad Sci U S A 104, 14489–14494.

Kuijpers, M., Kochlamazashvili, G., Stumpf, A., Puchkov, D., Swaminathan, A., Lucht, M.T., Krause, E., Maritzen, T., Schmitz, D., and Haucke, V. (2021). Neuronal Autophagy Regulates Presynaptic Neurotransmission by Controlling the Axonal Endoplasmic Reticulum. Neuron 109, 299–313.e299.

Lachance, V., Wang, Q., Sweet, E., Choi, I., Cai, C.Z., Zhuang, X.X., Zhang, Y., Jiang, J.L., Blitzer, R.D., Bozdagi-Gunal, O., et al. (2019). Autophagy protein NRBF2 has reduced expression in Alzheimer’s brains and modulates memory and amyloid-beta homeostasis in mice. Mol Neurodegener 14, 43.

Le Guerroué, F., Eck, F., Jung, J., Starzetz, T., Mittelbronn, M., Kaulich, M., and Behrends, C. (2017). Autophagosomal Content Profiling Reveals an LC3C-Dependent Piecemeal Mitophagy Pathway. Mol Cell 68, 786–796.e786.

Lee, J.H., Yang, D.S., Goulbourne, C.N., Im, E., Stavrides, P., Pensalfini, A., Chan, H., Bouchet-Marquis, C., Bleiwas, C., Berg, M.J., et al. (2022). Faulty autolysosome acidification in Alzheimer’s disease mouse models induces autophagic build-up of Abeta in neurons, yielding senile plaques. Nat Neurosci 25, 688–701.

Lee, S.J., Lodder, B., Chen, Y., Patriarchi, T., Tian, L., and Sabatini, B.L. (2021). Cell-type-specific asynchronous modulation of PKA by dopamine in learning. Nature 590, 451–456.

Longo, P.A., Kavran, J.M., Kim, M.S., and Leahy, D.J. (2013). Transient mammalian cell transfection with polyethylenimine (PEI). Methods Enzymol 529, 227–240.

Love, M.I., Huber, W., and Anders, S. (2014). Moderated estimation of fold change and dispersion for RNA-seq data with DESeq2. Genome Biol 15, 550.

Maday, S., and Holzbaur, E.L. (2014). Autophagosome biogenesis in primary neurons follows an ordered and spatially regulated pathway. Dev Cell 30, 71–85.

Mancias, J.D., Wang, X., Gygi, S.P., Harper, J.W., and Kimmelman, A.C. (2014). Quantitative proteomics identifies NCOA4 as the cargo receptor mediating ferritinophagy. Nature 509, 105–109.

Marro, S., and Yang, N. (2014). Transdifferentiation of mouse fibroblasts and hepatocytes to functional neurons. Methods Mol Biol 1150, 237–246.

Matsunaga, K., Saitoh, T., Tabata, K., Omori, H., Satoh, T., Kurotori, N., Maejima, I., Shirahama-Noda, K., Ichimura, T., Isobe, T., et al. (2009). Two Beclin 1-binding proteins, Atg14L and Rubicon, reciprocally regulate autophagy at different stages. Nat Cell Biol 11, 385–396.

Matsuoka, Y., Hughes, C.A., and Bennett, V. (1996). Adducin regulation. Definition of the calmodulin-binding domain and sites of phosphorylation by protein kinases A and C. J Biol Chem 271, 25157–25166.

McKnight, N.C., Zhong, Y., Wold, M.S., Gong, S., Phillips, G.R., Dou, Z., Zhao, Y., Heintz, N., Zong, W.X., and Yue, Z. (2014). Beclin 1 is required for neuron viability and regulates endosome pathways via the UVRAG-VPS34 complex. PLoS Genet 10, e1004626.

Menzies, F.M., Fleming, A., Caricasole, A., Bento, C.F., Andrews, S.P., Ashkenazi, A., Fullgrabe, J., Jackson, A., Jimenez Sanchez, M., Karabiyik, C., et al. (2017). Autophagy and Neurodegeneration: Pathogenic Mechanisms and Therapeutic Opportunities. Neuron 93, 1015–1034.

Mizushima, N. (2009). Methods for monitoring autophagy using GFP-LC3 transgenic mice. Methods Enzymol 452, 13–23.

Mizushima, N., and Komatsu, M. (2011). Autophagy: renovation of cells and tissues. Cell 147, 728–741.

Mizushima, N., Yamamoto, A., Matsui, M., Yoshimori, T., and Ohsumi, Y. (2004). In vivo analysis of autophagy in response to nutrient starvation using transgenic mice expressing a fluorescent autophagosome marker. Mol Biol Cell 15, 1101–1111.

Niu, M., Cho, J.H., Kodali, K., Pagala, V., High, A.A., Wang, H., Wu, Z., Li, Y., Bi, W., Zhang, H., et al. (2017). Extensive Peptide Fractionation and y1 Ion-Based Interference Detection Method for Enabling Accurate Quantification by Isobaric Labeling and Mass Spectrometry. Anal Chem 89, 2956–2963.

Nixon, R.A. (2013). The role of autophagy in neurodegenerative disease. Nat Med 19, 983–997.

Ordureau, A., Kraus, F., Zhang, J., An, H., Park, S., Ahfeldt, T., Paulo, J.A., and Harper, J.W. (2021). Temporal proteomics during neurogenesis reveals large-scale proteome and organelle remodeling via selective autophagy. Mol Cell 81, 5082–5098 e5011.

Øverbye, A., Fengsrud, M., and Seglen, P.O. (2007). Proteomic analysis of membrane-associated proteins from rat liver autophagosomes. Autophagy 3, 300–322.

Palmer, D.S., Howrigan, D.P., Chapman, S.B., Adolfsson, R., Bass, N., Blackwood, D., Boks, M.P.M., Chen, C.Y., Churchhouse, C., Corvin, A.P., et al. (2022). Exome sequencing in bipolar disorder identifies AKAP11 as a risk gene shared with schizophrenia. Nat Genet 54, 541–547.

Peng, J., Elias, J.E., Thoreen, C.C., Licklider, L.J., and Gygi, S.P. (2003). Evaluation of multidimensional chromatography coupled with tandem mass spectrometry (LC/LC-MS/MS) for large-scale protein analysis: the yeast proteome. J Proteome Res 2, 43–50.

Roney, J.C., Li, S., Farfel-Becker, T., Huang, N., Sun, T., Xie, Y., Cheng, X.T., Lin, M.Y., Platt, F.M., and Sheng, Z.H. (2021). Lipid-mediated motor-adaptor sequestration impairs axonal lysosome delivery leading to autophagic stress and dystrophy in Niemann-Pick type C. Dev Cell 56, 1452–1468.e1458.

Sahoo, S.K., and Kim, D.H. (2010). Characterization of calumenin in mouse heart. BMB Rep 43, 158–163.

Schmitt, D., Bozkurt, S., Henning-Domres, P., Huesmann, H., Eimer, S., Bindila, L., Tascher, G., Münch, C., Behl, C., and Kern, A. (2021). Protein content and lipid profiling of isolated native autophagosomes. bioRxiv, 2021.2004.2016.440117.

Sharoar, M.G., Hu, X., Ma, X.M., Zhu, X., and Yan, R. (2019). Sequential formation of different layers of dystrophic neurites in Alzheimer’s brains. Mol Psychiatry 24, 1369–1382.

Shen, B., Zheng, P., Qian, N., Chen, Q., Zhou, X., Hu, J., Chen, J., and Teng, J. (2019). Calumenin-1 Interacts with Climp63 to Cooperatively Determine the Luminal Width and Distribution of Endoplasmic Reticulum Sheets. iScience 22, 70–80.

Smith, M.D., Harley, M.E., Kemp, A.J., Wills, J., Lee, M., Arends, M., von Kriegsheim, A., Behrends, C., and Wilkinson, S. (2018). CCPG1 Is a Non-canonical Autophagy Cargo Receptor Essential for ER-Phagy and Pancreatic ER Proteostasis. Dev Cell 44, 217–232 e211.

Tan, H., Wu, Z., Wang, H., Bai, B., Li, Y., Wang, X., Zhai, B., Beach, T.G., and Peng, J. (2015). Refined phosphopeptide enrichment by phosphate additive and the analysis of human brain phosphoproteome. Proteomics 15, 500–507.

Tang, G., Gudsnuk, K., Kuo, S.H., Cotrina, M.L., Rosoklija, G., Sosunov, A., Sonders, M.S., Kanter, E., Castagna, C., Yamamoto, A., et al. (2014). Loss of mTOR-dependent macroautophagy causes autistic-like synaptic pruning deficits. Neuron 83, 1131–1143.

Tanida, I., Yamaji, T., Ueno, T., Ishiura, S., Kominami, E., and Hanada, K. (2008). Consideration about negative controls for LC3 and expression vectors for four colored fluorescent protein-LC3 negative controls. Autophagy 4, 131–134.

Tian, R., Gachechiladze, M.A., Ludwig, C.H., Laurie, M.T., Hong, J.Y., Nathaniel, D., Prabhu, A.V., Fernandopulle, M.S., Patel, R., Abshari, M., et al. (2019). CRISPR Interference-Based Platform for Multimodal Genetic Screens in Human iPSC-Derived Neurons. Neuron 104, 239–255 e212.

Vargas, J.N.S., Hamasaki, M., Kawabata, T., Youle, R.J., and Yoshimori, T. (2022). The mechanisms and roles of selective autophagy in mammals. Nat Rev Mol Cell Biol.

Vijayan, V., and Verstreken, P. (2017). Autophagy in the presynaptic compartment in health and disease. J Cell Biol 216, 1895–1906.

Wang, Q.J., Ding, Y., Kohtz, D.S., Mizushima, N., Cristea, I.M., Rout, M.P., Chait, B.T., Zhong, Y., Heintz, N., and Yue, Z. (2006). Induction of autophagy in axonal dystrophy and degeneration. J Neurosci 26, 8057–8068.

Wang, X., Li, Y., Wu, Z., Wang, H., Tan, H., and Peng, J. (2014). JUMP: a tag-based database search tool for peptide identification with high sensitivity and accuracy. Mol Cell Proteomics 13, 3663–3673.

Wang, Z., Yu, K., Tan, H., Wu, Z., Cho, J.H., Han, X., Sun, H., Beach, T.G., and Peng, J. (2020). 27-Plex Tandem Mass Tag Mass Spectrometry for Profiling Brain Proteome in Alzheimer’s Disease. Anal Chem 92, 7162–7170.

Yamamoto, A., and Yue, Z. (2014). Autophagy and its normal and pathogenic states in the brain. Annu Rev Neurosci 37, 55–78.

Zellner, S., Nalbach, K., and Behrends, C. (2021a). Autophagosome content profiling using proximity biotinylation proteomics coupled to protease digestion in mammalian cells. STAR Protoc 2, 100506.

Zellner, S., Schifferer, M., and Behrends, C. (2021b). Systematically defining selective autophagy receptor-specific cargo using autophagosome content profiling. Mol Cell 81, 1337–1354.e1338.

Zhang, Y., Pak, C., Han, Y., Ahlenius, H., Zhang, Z., Chanda, S., Marro, S., Patzke, C., Acuna, C., Covy, J., et al. (2013). Rapid single-step induction of functional neurons from human pluripotent stem cells. Neuron 78, 785–798.

Zhong, H., Sia, G.M., Sato, T.R., Gray, N.W., Mao, T., Khuchua, Z., Huganir, R.L., and Svoboda, K. (2009a). Subcellular dynamics of type II PKA in neurons. Neuron 62, 363–374.

Zhong, Y., Wang, Q.J., Li, X., Yan, Y., Backer, J.M., Chait, B.T., Heintz, N., and Yue, Z. (2009b). Distinct regulation of autophagic activity by Atg14L and Rubicon associated with Beclin 1-phosphatidylinositol-3-kinase complex. Nat Cell Biol 11, 468–476.

